# Latitudinal trends in genetic diversity and distinctiveness of *Quercus robur* rear edge forest remnants call for new conservation priorities

**DOI:** 10.1101/2023.10.30.564819

**Authors:** Camilla Avanzi, Francesca Bagnoli, Edoardo Romiti, Ilaria Spanu, Yoshiaki Tsuda, Elia Vajana, Giovanni Giuseppe Vendramin, Andrea Piotti

## Abstract

Knowledge of the spatial distribution of intraspecific genetic variation is essential for planning conservation actions, designing networks of protected areas, and informing possible assisted migration strategies. Although the Italian peninsula harbours unique genetic variation as a legacy of Quaternary migrations, only sporadic genetic information is available for forest tree species.

Here, we present the first geographically comprehensive genetic characterisation of *Quercus robur*, an iconic broadleaved species of European floodplain forests, in an area which acted as a primary glacial refugium for the species. 745 individuals from 25 populations were sampled and genotyped with 16 microsatellite markers. Their genetic structure was assessed through various metrics of diversity and distinctiveness, as well as by Bayesian clustering and multivariate methods. The demographic history of inferred gene pools was evaluated through Approximate Bayesian Computation analysis.

Genetic distinctiveness showed a decline with increasing latitude, while allelic richness reached its peak in central Italy. A south-to-north trend in the complexity of the genetic structure was observed, with peninsular Italy being characterised by intermingled gene pools in contrast to the relative homogeneity exhibited by northern populations. Demographic inference indicated that the southern gene pool has been genetically isolated since the penultimate interglacial, whereas populations from central Italy persisted locally in a mosaic of small refugia during the last glaciation.

Our results brought out the complexity of the genetic structure of forest trees’ populations in southern Europe. At least three *Q. robur* refugia contributed to the genetic layout of extant populations after the last glacial period, but refugial areas were probably even more numerous in central Italy. Such a detailed characterization sheds new light on the priorities to be established for the conservation of highly fragmented *Q. robur* populations in an area rich in diversified genetic lineages.

## Introduction

The DNA-encoded heritable variation within and between populations of a species (i.e. intraspecific genetic diversity) provides species with the ability to adapt and evolve (Hoban et al. 2020) and influences ecological processes, with significant effects on community structure, primary production, nutrient cycling, and ecosystem resilience (Hughes et al. 2008). In the context of the current global climatic crisis, adequate levels of genetic diversity are the essential prerequisite for species to keep pace with rapidly changing environmental conditions and the spread of new pests and diseases (Sgrò et al. 2011), as well as for ecosystems to keep providing ecological functions and services (Des Roches et al. 2021).

Despite broad scientific *consensus* on the urgency of conserving intraspecific genetic diversity, and hence adaptive potential (DeWoody et al. 2021), conservation strategies often provide vague guidance on how to translate this goal into practical action, particularly at national and subnational levels (Pierson et al. 2016; Cook and Sgrò 2017; Laikre et al. 2020). Hoban et al. (2020) proposed three pragmatic indicators for genetic monitoring, based on the number of populations with an effective population size (*Ne*) above the standard (but controversial) threshold of 500, the proportion of populations conserved for each species and, finally, the number of populations in which genetic diversity is assessed (and tracked) using molecular markers. Indeed, direct knowledge of the spatial distribution of intraspecific genetic diversity and the degree of genetic connectivity among populations is particularly useful for informing translocation strategies and/or designing optimal networks of protected areas (Aitken and Whitlock 2013; Fady et al. 2016; Andrello et al. 2022). Despite the fact that range-wide surveys of the genetic variation of European forest tree species have been carried out for a long time (e.g. Petit et al. 2003; Vendramin et al. 2008; Postolache et al. 2021; Milesi et al. 2023), southern Europe has frequently been under-sampled, despite its disproportionate importance as a long-term repository of genetic diversity (Hampe and Petit 2005). Southern Europe likely hosted a mosaic of refugial areas for tree species during the cold phases of the Quaternary (Tzedakis et al. 1997), resulting in southern European populations often exhibiting complex spatial patterns in the distribution of genetic diversity (e.g. Piotti et al. 2017; Scotti-Saintagne et al. 2021). Robust sampling within putative refugial areas is essential to quantitatively describe such complexity (Piotti et al. 2017) and, in turn, to design effective conservation strategies of extant forest genetic resources (de Vries et al. 2015).

Pedunculate oak (*Quercus robur* L.) is an iconic broadleaved tree species key for the European lowland ecosystems, ranging from Mediterranean peninsulas to Scandinavia, from Scotland to the Ural and Caucasus mountain ranges (Eaton et al. 2016). Although *Q. robur* may appear to be distributed across a continuous range according to the available distribution maps (Caudullo et al. 2017), it regionally suffers from a high degree of fragmentation, as lowland forests have been increasingly replaced by agricultural and urban landscapes in large parts of the species’ distribution (Petit et al. 2002c; Vakkari et al. 2006; Belletti et al. 2007; Agnoletti et al. 2018; Degen et al. 2021). In recent decades, and particularly in rear edge populations, fragmentation has been exacerbated by episodes of decline and mortality, fuelled by a complex interplay of changing biotic and abiotic conditions (Gibbs 1997; Ragazzi et al. 2000; Thomas et al. 2002; de Sampaio e Paiva Camilo-Alves et al. 2013; Stojanović et al. 2015; Gathercole et al. 2021). In this context, a comprehensive and accurate study of the genetic layout of *Q. robur* populations at the rear edge of the species’ distribution within a putatively multi-refugial area, such as the Italian peninsula, is still lacking. To our knowledge, despite a few studies at regional scale (Belletti et al. 2005; Ducci et al. 2007), only Fineschi et al. (2002) investigated the genetic diversity of *Q. robur* populations throughout Italy, as part of a larger study of chloroplast DNA variation of European white oaks (Petit et al. 2002a, b). However, this study relied on a very limited *Q. robur* sample size (64 trees), with only a few samples from peninsular Italy. Nevertheless, there is evidence that this area hosted multiple glacial refugia for several forest tree species (Magri et al. 2015) and that it was relevant for post-glacial demographic history of white oaks in Europe (Petit et al. 2002a). Sparse genetic information and the impossibility to distinguish *Quercus* pollen at the species level prevent testing specific hypotheses on *Q. robur* recent demographic dynamics and call for a deep investigation on the genetic structure of the species, and the major events influencing it, in this area.

Our work is the first attempt to comprehensively characterise the forest genetic resources of pedunculate oak throughout Italy. To this end, 25 populations were carefully selected, sampled and genotyped with 16 microsatellite markers. The specific aims of this study were to: i) assess their levels of genetic diversity and distinctiveness, ii) quantify the contribution of each population to the total allelic diversity and its components within and between populations, iii) identify which populations have limited effective population size and show genetic signatures of bottlenecks, iv) decipher the extant genetic structure of the species along the Italian peninsula, and v) date the divergence of extant gene pools. As peninsular Italy is characterised by greater topographic heterogeneity than the Po Valley, and it hosted multiple forest trees’ refugial areas, genetic diversity and distinctiveness, as well as the complexity of genetic structure, are expected to decrease with increasing latitude. Uncovering hotspots of genetic diversity and distinctiveness, as well as the existence of potential gene flow constraints among oak forest remnants, will be essential to inform management and conservation actions aimed at mitigating the negative effects of fragmentation genetic drift in such an economically and ecologically important tree species.

## Materials and methods

### Sampling activities and genotyping

Twenty-five populations of *Quercus robur* L. were sampled throughout Italy (Figure 1). In each population, with few exceptions due to extremely small natural stands (Table S1), leaves were collected from ∼30 individuals spaced at least 25 meters to avoid sampling related trees, reaching a total of 745 sampled trees. Each individual was tagged and georeferenced using a GPS tracker. Fresh leaves were dried in silica gel and stored at -20 °C until DNA extraction. To be sure that we had not mistakenly sampled individuals of other white oak species, we also collected leaves from 46, 26 and 8 trees of *Q. petraea*, *Q. pubescens* and *Q. frainetto*, respectively (Figure S1a) for successive species discrimination based on the same set of molecular markers.

**Figure 1.**
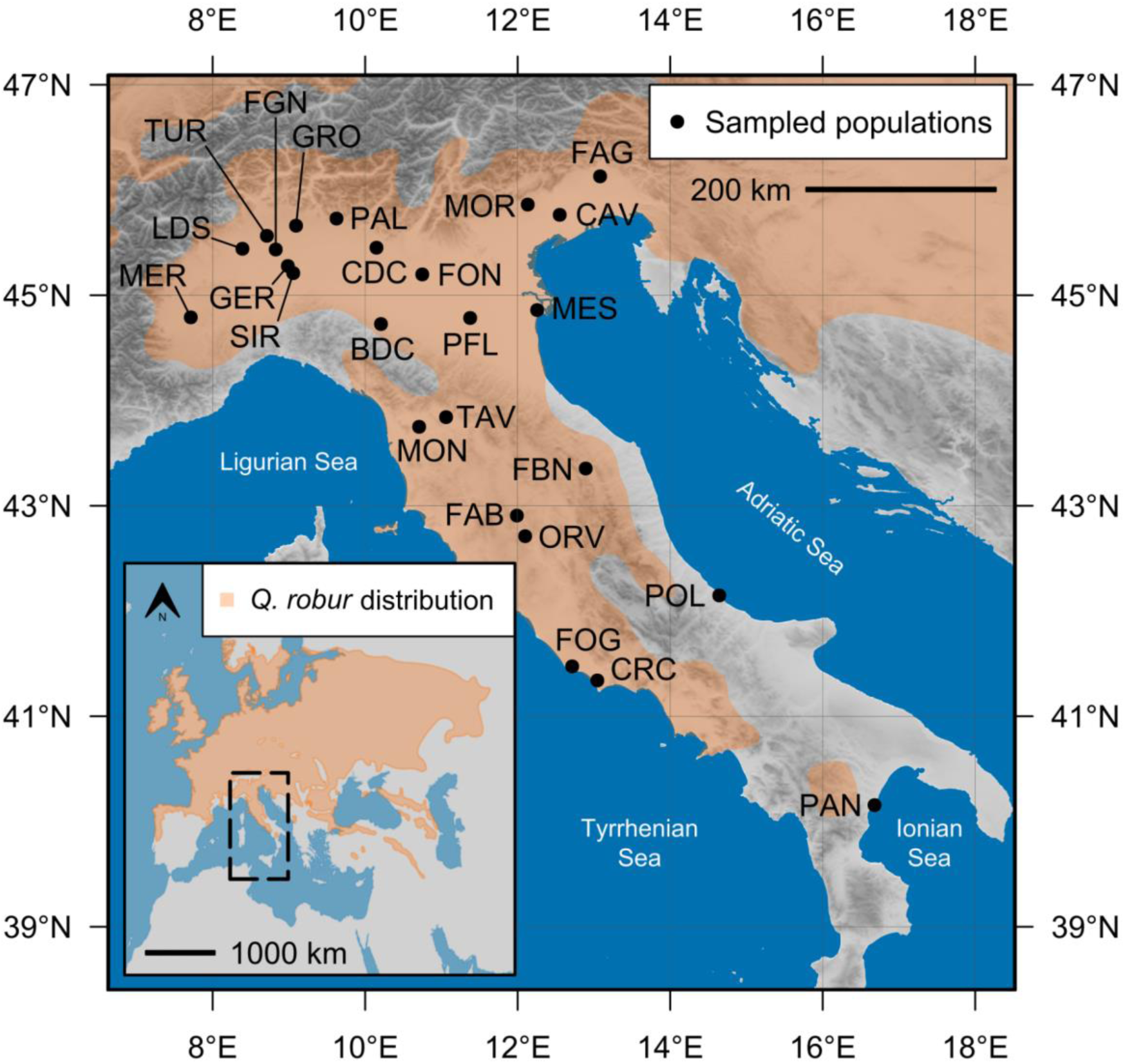
Map of *Quercus robur* populations sampled for the present study. The inset represents the distribution of the species in Europe according to Caudullo et al. (2017). The distribution map of the species strongly overestimates, at least for Italy, the extent and continuity of *Q. robur*-dominated formations, whose distribution is in fact largely fragmented (Belletti et al. 2007).

DNA was extracted using the NucleoSpin 96 Plant II Kit (Macherey-Nagel, Germany), following the manufacturer’s instructions. Approximately 30 mg of frozen leaves and a 3-mm tungsten bead were added to each well of 96-well plates for grinding leaf material. Then, plates were frozen at -80 °C before two cycles of 30-second disruption at 27 Hz using a Mixer Mill MM300 (Retsch, Germany). All individuals were genotyped with 16 nuclear microsatellite markers (SSRs) (PIE020, PIE223, PIE242, PIE102, PIE239, PIE227, PIE271, PIE267, PIE215: Durand et al. 2010; MsQ13: Dow et al. 1995; QrZAG7, QrZAG112, QrZAG20, QrZAG96:

Kampfer et al. 1998; QpZAG15, QpZAG110: Steinkellner et al. 1997). All 16 SSRs were arranged into a 9- and 7-plex by slightly modifying the 12-plex and 8-plex developed by Guichoux et al. (2011). Multiplex reactions were carried out with the Type-it Microsatellite PCR Kit (Qiagen, Germany). The final volume of the PCR reactions was optimised to 6 μl. The PCR mix was composed of 3 μl Type-it Multiple PCR Master Mix, 0.6 μl of primer premix, 1.4 μl of H_2_O and 1 μl of DNA. The cycling conditions indicated by Guichoux et al. (2011) were used for the 9-plex, while they were slightly changed for the 7-plex, with an initial step at 95°C for 5 min, followed by 32 cycles at 95°C for 30 s, 56°C for 90 s and 72°C for 45 s, and a final 30 min extension step at 60°C. PCR products were run on an ABI 3500 automatic sequencer (Applied Biosystems, USA), with LIZ-500 (ThermoFisher Scientific) as internal size standard. The obtained results were sized using GeneMarker (Softgenetics).

### Quality control of the genetic dataset

In some populations of the present study, *Quercus robur* was in close proximity with other related oak species (i.e. *Q. petraea*, *Q. pubescens*, *Q. frainetto*). Although some ‘easy-to-use’ discriminant morphometric traits were used during fieldwork to identify and sample *Q. robur* trees, molecular markers were subsequently used to further exclude the presence of individuals ‘misclassified’ as *Q. robur*, which would bias estimates of genetic diversity and structure. Our set of SSRs was indeed proven to discriminate well European white oaks (Guichoux et al. 2011). Such a check was done by applying the Bayesian clustering approach implemented in STRUCTURE (Pritchard et al. 2000), with sampling locations as prior information (Hubisz et al. 2009). Thirty independent runs were run for each value of *K* (the number of putative gene pools), ranging from two to four (i.e. the number of white oak species). Each run consisted of 10,000 burn-in iterations and 50,000 sampling iterations. Runs were aligned and averaged using the software CLUMPAK (Kopelman et al. 2015). Individuals with a probability of belonging to one of the other white oak species >0.8 were considered ’misclassified’ and excluded from any further analysis. The Bayesian clustering approach was complemented by a multivariate analysis, i.e. a Principal Coordinate Analysis (PCoA) performed on the matrix of pairwise mean among-population genetic distances (Smouse and Peakall 1999) using GenAlEx (Peakall and Smouse 2012).

The presence of null alleles and genotyping failures in the final dataset (i.e. the dataset without ‘misclassified’ individuals) was assessed at the population level by using INest v2.2 (Chybicki and Burczyk 2009). The model was run using a Gibbs sampler based on 200,000 iterations, with a burn-in of 20,000 and retaining one iteration every 200.

To ensure the neutrality of the marker set, first 10,000 coalescent simulations were performed in the Arlequin v3.5.2 environment (Excoffier and Lischer 2010) to get the p-values of locus-specific F-statistic conditioned on the observed levels of heterozygosity (Beaumont and Nichols 1996; Excoffier et al. 2008). A non-hierarchical finite island model was used, with all populations pooled into a single group. Secondly, the hierarchical Bayesian approach implemented in BayeScan v2.1 (Foll and Gaggiotti 2008) was used to estimate the posterior probabilities of two alternative models, one including the effect of selection and one excluding it. The ratio of the posterior probabilities of the two models can be translated, for each locus, into a different level of selection intensity according to Jeffreys’ interpretation. BayeScan was run on demographically stable populations identified based on the results of a bottleneck analysis carried out with INest (for details see the ‘Demographic parameters’ paragraph) with 1000 pilot runs of 10,000 iterations each, and a burn-in of 500,000, a sample size of 10,000 and a thinning interval of 20. Prior odds for the selection model were set to 100.

### Genetic structure

The Bayesian clustering approach implemented in STRUCTURE was used to identify the most likely number of gene pools (*K*). The analysis was based on the LOCPRIOR model (Hubisz et al. 2009), as well as on the correlated allele frequencies model (Falush et al. 2003). Thirty independent runs were performed for each value of *K*, ranging from 1 to 10. Each run consisted of 10,000 burn-in iterations and 50,000 data collection iterations. These settings for the total length of the MCMC were sufficient to obtain reliable genetic structures by comparing the results of initial runs with those involving longer chains. Different runs for the same *K* were then averaged using the software CLUMPAK. The most likely grouping scenarios were evaluated by using Δ*K* statistic (Evanno et al. 2005), as well as by a visual assessment of the log-likelihood trend across the various tested scenarios. Both Δ*K* and log-likelihood values were calculated by using the web-based software StructureSelector (Li and Liu 2018).

The genetic structure of Italian oak populations was also investigated with a multivariate approach, running a PCoA on the matrix of pairwise among-population genetic distance (Smouse and Peakall 1999) with GenAlEx. Pairwise *G*_ST_ values (Meirmans and Hedrick 2011) of genetic differentiation among populations were also calculated with GenAlEx, testing their significance by 999 permutations.

### Genetic diversity and distinctiveness

Number of alleles (*Na*), as well as the observed (*H*_O_) and expected heterozygosity (*H*_E_), were calculated for each population using GenAlEx. The inbreeding coefficient (*F*_IS_) of each population was assessed by using INEST. Allelic richness (*Ar*) and private allelic richness (*PAr*) were calculated by using HP-RARE (Kalinowski 2005). Rarefaction was applied based on a minimum sample size of 44 genes. The contribution of each population to the total allelic diversity (*A*_T_), together with its within-subpopulation (*A*_S_) and among-subpopulation (*D*_A_) components, was assessed using METAPOP2 (López-Cortegano et al. 2019).

The genetic distinctiveness of each population was estimated using a population-specific index of differentiation (*F*_ST_) between the target population and a common migrant gene pool from which it differs in varying degrees, using BayeScan v2.1. *F*_ST_ coefficients were estimated through 20 pilot runs of 10,000 iterations each, followed by a final MCMC made up of a burn-in of 50,000 and further 50,000 iterations with a thinning of 20.

Linear regression was used to explore the relationship between genetic parameters (i.e. Na, *H*_O_, *H*_E_, *F*_IS_, *Ar, PAr, A*_T_, *A*_S_, *D*_A_, *F*_ST_) and latitude. To account for non-linear relationships, a quadratic term of latitude was tested in the models. Global *P*-values were adjusted for multiple testing using the Bonferroni correction. Regression models and multiple testing correction were performed using the *lm* and *p.adjust* R functions from the *stats* package (R Core Team 2023)

### Demographic parameters

Effective population size (*Ne*) was estimated using the linkage disequilibrium method implemented in NeEstimator v2.1 (Do et al. 2014). In forest tree species, due to their peculiar life-history traits, reliable estimates of *Ne* are expected only for isolated and small populations (typically less than 500 reproductive trees) (Santos-del-Blanco et al. 2022). In our dataset, the populations that met such requirements are shown in Table S1.

The occurrence of past genetic bottlenecks within each population was evaluated by searching both for an excess of heterozygosity (*H*; Cornuet and Luikart 1996) and a deficiency in the ratio between the number of alleles and the range of allele size (MR; Garza and Williamson 2001) with INEST. The values of *H* and MR expected at equilibrium (*H*_eq_ and MR_eq_) were computed with coalescent simulations, assuming a two-phase mutation model with a proportion and average size of multi-step mutations of 0.22 and 3.1, respectively, as recommended by Peery et al. (2012). *H*_eq_ and MR_eq_ were then tested against observed *H* and MR values using a one-tailed Wilcoxon signed-rank test with 1,000,000 permutations (no normality assumption required).

### Approximate Bayesian Computation to trace back the divergence among oak gene pools

To infer the past demographic history of Italian *Quercus robur* populations, the Approximate Bayesian Computation (ABC) method implemented in the DIYABC v2.1 software (Cornuet et al. 2008, 2014) was applied. To limit the number of tested scenarios, the genetic information obtained at *K*=3 from the Bayesian clustering analysis was used to guide the ABC analysis. Therefore, three populations were defined: NI (northern Italy), CI (central Italy), and SI (southern Italy) (see Results, Figure 3b). NI and CI were assembled by randomly selecting 60 individuals from the populations assigned to the relative gene pool with a membership ≥0.7, while SI consisted of the entire PAN population (61 individuals). After some pilot runs to determine which was the ancestral population (Appendix 1) and the topology of the split scenarios (results not shown), four demographic scenarios were compared (Figure 2a). In all tested scenarios, t# refers to the time scale measured in generation time, N# stands for the effective population sizes of the corresponding gene pool (i.e. NI, CI, SI, ancestral populations ‘a’, and changes in the effective population seize ‘b’), and r# represents the admixture ratio. In principle, DIYABC does not consider the hypothesis of post-split gene flow within scenarios, but a recent study modified DIYABC to include symmetric admixture between two populations (Chapuis et al. 2020). Kitamura et al. (2022) extended this approach to model directional admixture (e.g. from NI to CI and from CI to NI), which could be considered analogous to gene flow. Thus, the parameter ’r#’ related to the admixture rate between two generic populations *a* and *b* can be considered as the relative amount of gene flow at time ’t#’, while ’1-r#’ represents the value of the relative amount of gene flow within the reference parent population.

**Figure 2.**
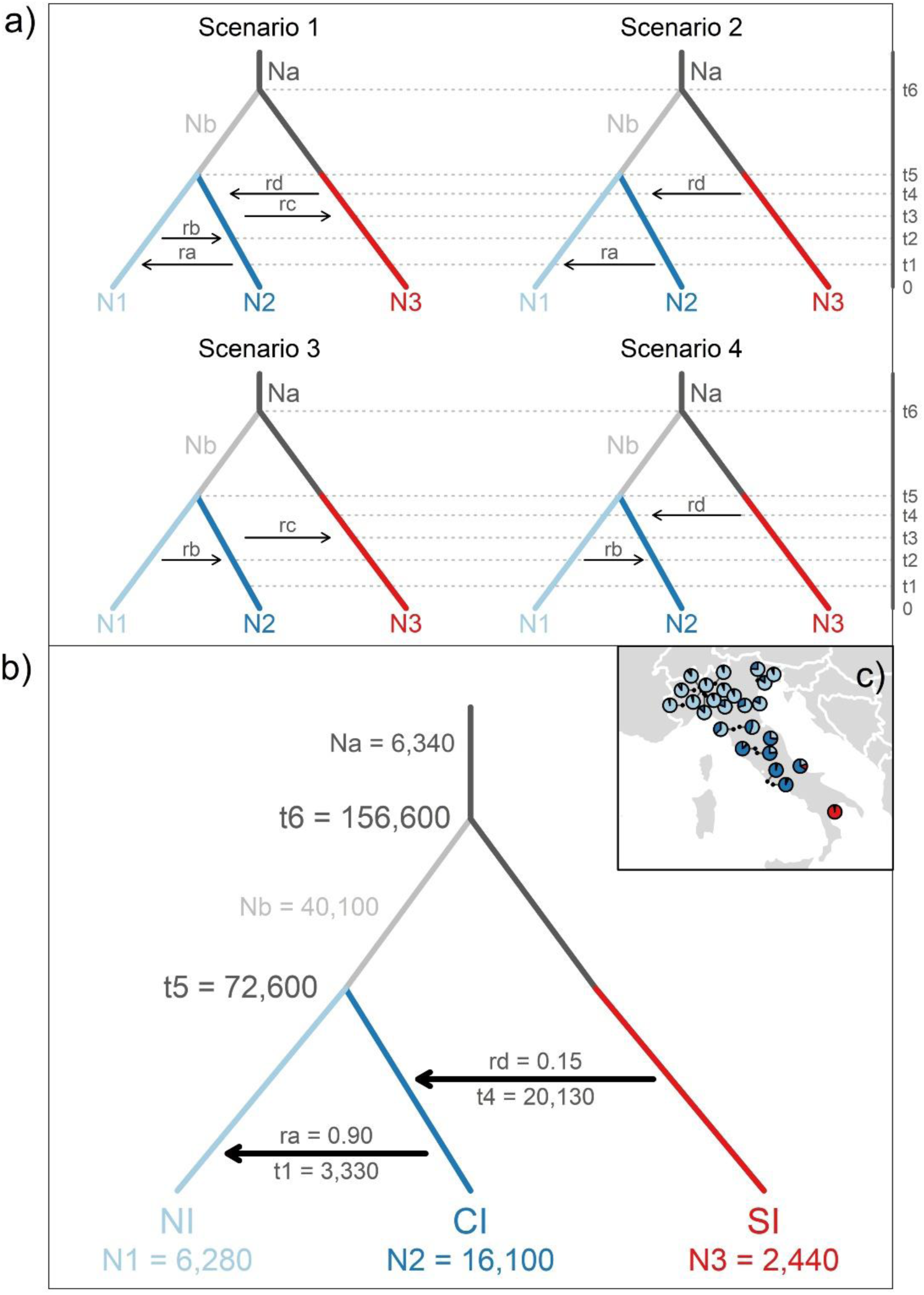
a) Four demographic models tested by using the Approximate Bayesian Computation (ABC) approach implemented in DIYABC. N1, N2 and N3 represent the effective population size of NI, CI and SI, that are the three gene pools resulting from the Bayesian clustering analysis and represent, respectively, individuals from northern Italian populations, central Italian populations and the southernmost stand of PAN. Na and Nb represent the effective population size of the ancestral populations ‘a’ and ‘b’. t# = time-scaled by generation time; r = admixture ratio. b) Median values of posterior parameters for *Scenario 2*, which turned out to be the most likely demographic model based on posterior probabilities. c) Visualization of the geographic positions of the three gene pools, based on the results of the Bayesian clustering approach at *K* = 3 (see Figure 3 for more details).

**Figure 3.**
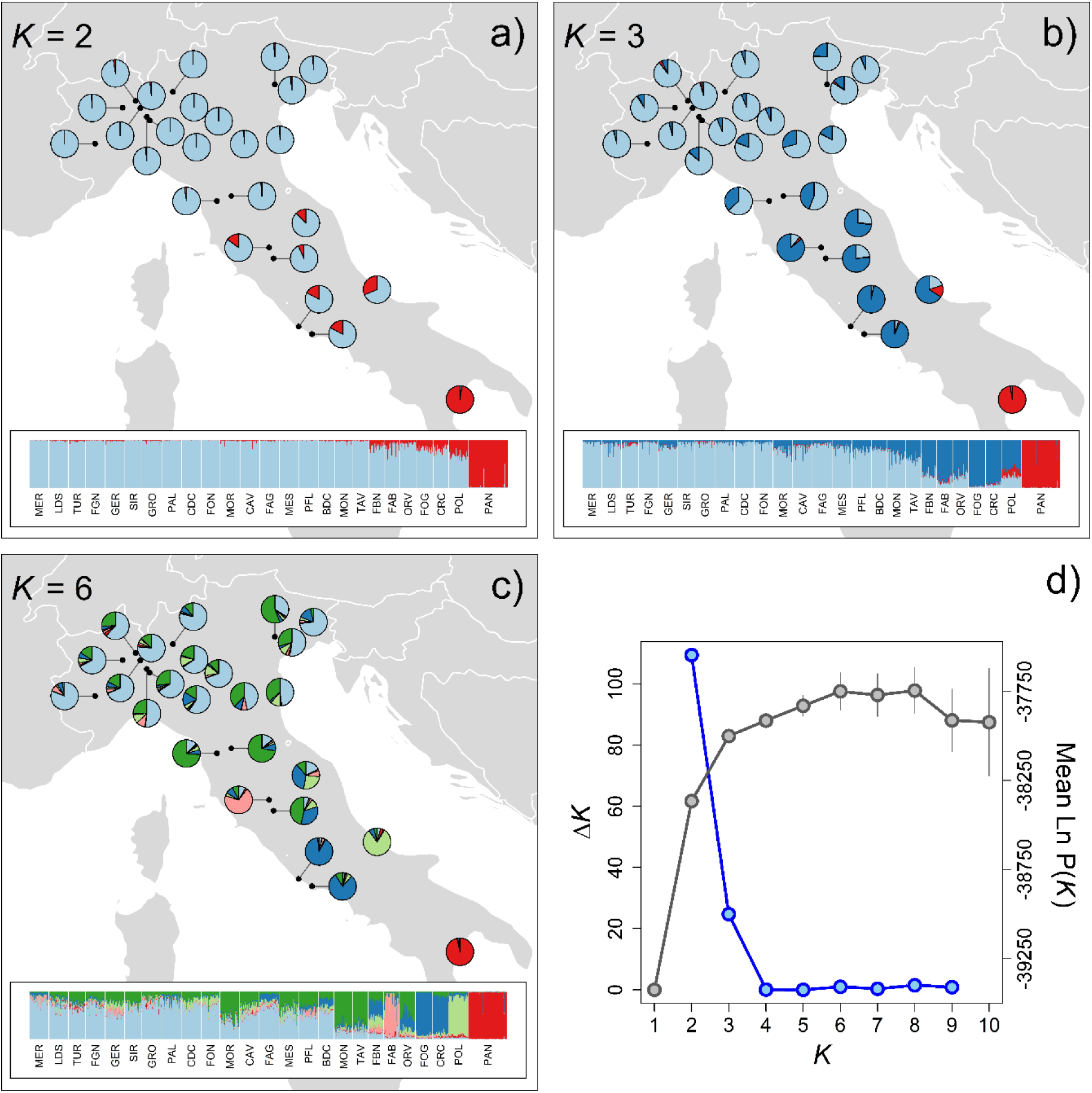
Results of the Bayesian clustering analysis carried out with STRUCTURE. For each population, average ancestry coefficients at a) *K* = 2, b) *K* = 3, and c) *K* = 6 are shown. For each structuring level, individual ancestry coefficients are also shown in the barplot at the bottom of each panel. Evanno et al. (2005)’s Δ*K* and log-likelihood values used to evaluate the most likely grouping scenarios are presented in panel d).

All scenarios were based on a *hierarchical split model* which, as suggested by the results of the pilot runs (Appendix 1), assumed that the SI gene pool could be traced back to the ancestral population (Na). The model also predicted a split between SI and the gene pools further north at t6, and a divergence between NI and CI at t5. At t5, changes in population size on both branches of the tree (Na and Nb) were also modelled. The differences between the four scenarios were based on patterns of ’gene flow’ (*sensu* Kitamura et al. 2022) modelled using a ’stepping stone’ approach, i.e. gene flow was only possible between geographically close gene pools (e.g. from SI to CI, but not from SI to NI):

- in *Scenario 1*, gene flow was bidirectional, from north to south (i.e. rb from NI to CI at t2 and rc from CI to SI at t3) and from south to north (i.e. rd from SI to CI at t4 and ra from CI to NI at t1);
- in *Scenario 2*, gene flow was unidirectional, from south to north (i.e. rd from SI to CI at t4 and ra from CI to NI at t1);
- in *Scenario 3*, gene flow was unidirectional, from north to south (i.e. rb from NI to CI at t2 and rc from CI to SI at t3);
- in *Scenario 4*, there was gene flow converging to central Italy, from north (i.e. rb from NI to CI at t2) and from south (i.e. rd from SI to CI at t4).

The Generalized Stepwise Mutation model (GSM; Estoup et al. 2002) with Single Nucleotide Indels (SNI) was used for all ABC simulations. The default priors were modified to obtain better posterior distributions based on the results of the pilot runs (Table S2). The minimum and maximum priors for SSR mutation rate were set to 1×10^-4^–1×10^-3^. The mean number of alleles, mean genetic diversity, mean allele size variance and mean Garza-Williamson’s M were used as summary statistics for individual populations. For each population pair, mean number of alleles, mean genetic diversity, mean allele size variance, *F*_ST_, and mean individual assignment likelihood were calculated. One million simulations were run for each scenario. After a total of four million simulations, the most likely scenario was assessed by comparing posterior probabilities, using logistic regression. Goodness-of-fit was estimated for each scenario by model checking using the Principal Component Analysis (PCA) approach implemented in DIYABC, which measures the discrepancy between simulated and observed data.

## Results

### Quality control of the genetic dataset

The Bayesian clustering approach used to detect potential ‘misclassified’ *Q. robur* individuals showed that *Q. petraea*, *Q. pubescens* and *Q. frainetto* trees clustered together at *K* = 2; at *K* = 3 the southern population of PAN emerged as a distinct gene pool, while at *K* = 4, central Italy populations showed distinctive genetic features (Figure S1c). *Q. petraea*, *Q. pubescens* and *Q. frainetto* individuals still clustered together at *K* = 3 and *K* = 4 but were discriminated against the second axis of the Principal Coordinate Analysis (PCoA) (Figure S1b). Based on STRUCTURE results at *K* = 4, 16 individuals were identified as ‘misclassified’ trees and were excluded from the final dataset. They belonged to four populations: FON (one single tree), MON (ten trees), FBN (two trees) and PAN (three trees) (see Table S1).

No marker showed consistent and/or significant issues related to genotyping failure and null alleles (Figure S2) and significant signals of selection based on the coalescent simulations performed within the Arlequin environment (Table S3). The Bayesian approach implemented in BayeScan showed a signature of balancing selection for QrZag112, with a *F*_ST_ value close to zero (Table S4). As the genetic structure and allelic richness with and without this marker was almost identical (results not shown), and the locus was particularly informative for detecting past demographic dynamics (it had the highest number of alleles), QrZag112 was retained in the final dataset.

### Genetic structure

According to Δ*K* statistic, the Bayesian clustering analysis revealed an optimal grouping at *K* = 2 (Figure 3d), which clearly distinguished a gene pool represented by the southernmost population of PAN from another gene pool represented by all the other Italian populations (Figure 3a). Mean pairwise *G*_ST_ value between PAN and all other populations was 0.044 (Table S5). Based on Δ*K*, the next strongest level of structuring was *K* = 3 (Figure 3d), which separated populations from central Italy from those from northern Italy (Figure 3b). Although Δ*K* was the highest when *K* = 2, the mean Ln P(*K*) increased with *K,* reaching its maximum at *K* = 6 (Figure 3d). According to this scenario, the complexity of the genetic structure in central Italy increased while populations from northern Italy mostly kept on clustering together (Figure 3c). Mean *r* values (averaged over runs for the same *K* value) were equal to 0.71, 0.45 and 0.54 when analysing two, three and six gene pools, respectively, indicating that the sampling locations were informative for unravelling the genetic structure. The results of the Principal Coordinate Analysis (PCoA) on the matrix of pairwise genetic distance among populations confirmed the genetic structure observed with the Bayesian clustering approach (Figure S3). The first three coordinates explained 60.3% of the total variance of the SSRs data. The southernmost population of PAN was well separated from all the others both on the first and second PCo axes. FAB population showed a clear separation from all the other populations on the third PCo axis. Pairwise indexes of genetic differentiation among populations (G_ST_) were reported in Table S5.

### Genetic diversity and distinctiveness

The 16 SSRs showed a total and mean number of alleles of 246 and 15.4, respectively. Estimates of population genetic diversity and distinctiveness are reported in Table 1. *Ar* showed a quadratic dependence on latitude (*lat*: F-value = 5.69, *P* = 0.026; *lat*^2^: F-value = 8.88, *P* = 0.007), with populations from central Italy showing the highest *Ar* values (Figure 4a). Genetic distinctiveness (population-specific *F*_ST_) had a negative relationship with latitude (*lat*: F-value = 53.65, *P* = 2.474×10^-7^; *lat*^2^: F-value = 16.86, *P* = 4.65×10^-4^; Figure 4b). Na, *PAr*, *H*_O_, *H*_E_, and *F*_IS_ showed no significant relationship with latitude (Figure S4).

**Table 1.** Genetic characteristics of sampled populations. *Na*: mean number of alleles, *Ar*: allelic richness, *PAr*: private allelic richness, *H*_O_: observed heterozygosity, *H*_E_: expected heterozygosity, F_IS_: inbreeding coefficient, *F*_ST_: plot-specific differentiation index according to Foll and Gaggiotti (2008), *Ne* and *Ne* CI_95%_: effective population size and its parametric 95% confidence interval.

| Population | *Na* | *Ar* | *PAr* | *H<sub>O</sub>* | *H<sub>E</sub>* | *F<sub>IS</sub>* | *F<sub>ST</sub>* | *Ne* | *Ne CI<sub>95%</sub>* |
| --- | --- | --- | --- | --- | --- | --- | --- | --- | --- |
| MER | 7.375 | 6.852 | 0.108 | 0.719 | 0.679 | 0.007 | 0.020 | 115 | 66-339 |
| LDS | 7.875 | 7.285 | 0.051 | 0.690 | 0.698 | 0.033 | 0.012 | 655 | 143-∞ |
| TUR | 7.688 | 7.350 | 0.008 | 0.676 | 0.691 | 0.026 | 0.013 | ∞ | 244-∞ |
| FGN | 7.875 | 7.241 | 0.059 | 0.679 | 0.690 | 0.014 | 0.010 | ∞ | 196-∞ |
| GER | 7.938 | 7.295 | 0.046 | 0.667 | 0.675 | 0.009 | 0.012 | 97 | 61-214 |
| SIR | 7.750 | 7.441 | 0.064 | 0.725 | 0.726 | 0.015 | 0.016 | 1330 | 139-∞ |
| GRO | 7.625 | 7.043 | 0.003 | 0.698 | 0.684 | 0.010 | 0.013 | ∞ | 382-∞ |
| PAL | 8.188 | 7.510 | 0.003 | 0.735 | 0.697 | 0.006 | 0.012 | 522 | 134-∞ |
| CDC | 7.375 | 6.841 | 0.122 | 0.671 | 0.684 | 0.021 | 0.023 | 106 | 61-315 |
| FON | 7.563 | 7.027 | 0.017 | 0.640 | 0.673 | 0.027 | 0.012 | 272 | 101-∞ |
| MOR | 7.625 | 7.062 | 0.086 | 0.721 | 0.683 | 0.004 | 0.025 | 46 | 35-65 |
| CAV | 7.938 | 7.319 | 0.016 | 0.688 | 0.696 | 0.016 | 0.008 | ∞ | 240-∞ |
| FAG | 7.375 | 6.801 | 0.026 | 0.654 | 0.654 | 0.011 | 0.025 | 1061 | 144-∞ |
| MES | 8.000 | 7.349 | 0.094 | 0.691 | 0.706 | 0.021 | 0.014 | 738 | 141-∞ |
| PFL | 7.750 | 7.190 | 0.047 | 0.685 | 0.694 | 0.019 | 0.014 | 285 | 107-∞ |
| BDC | 7.813 | 7.629 | 0.030 | 0.720 | 0.696 | 0.013 | 0.008 | ∞ | 256-∞ |
| MON | 7.688 | 7.193 | 0.073 | 0.682 | 0.684 | 0.016 | 0.025 | 325 | 104-∞ |
| TAV | 7.625 | 7.440 | 0.228 | 0.734 | 0.701 | 0.006 | 0.023 | 1348 | 144-∞ |
| FBN | 7.813 | 7.746 | 0.151 | 0.725 | 0.702 | 0.006 | 0.028 | 51 | 37-76 |
| FAB | 6.250 | 6.114 | 0.100 | 0.679 | 0.624 | 0.006 | 0.080 | 22 | 17-29 |
| ORV | 8.188 | 7.986 | 0.026 | 0.732 | 0.725 | 0.008 | 0.021 | ∞ | 184-∞ |
| FOG | 7.313 | 6.974 | 0.150 | 0.734 | 0.692 | 0.009 | 0.053 | 36 | 27-54 |
| CRC | 7.625 | 7.428 | 0.066 | 0.732 | 0.706 | 0.007 | 0.038 | 141 | 73-965 |
| POL | 7.000 | 6.522 | 0.115 | 0.679 | 0.666 | 0.011 | 0.055 | 60 | 41-102 |
| PAN | 6.375 | 5.352 | 0.151 | 0.655 | 0.651 | 0.013 | 0.165 | 92 | 67-135 |

**Figure 4.**
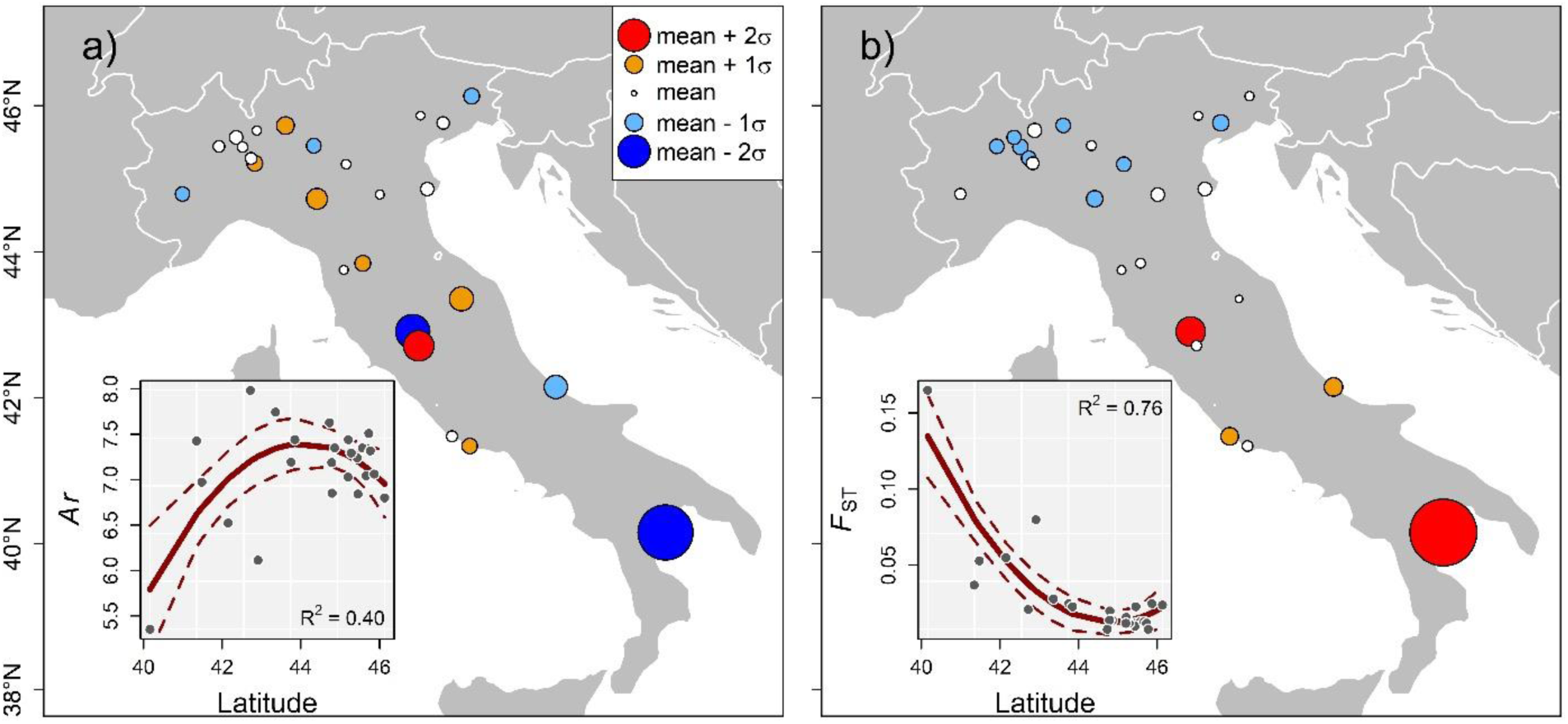
Estimates of (a) allelic richness (*Ar*), and (b) genetic distinctiveness (plot-specific *F*_ST_) in the 25 *Quercus robur* populations of the present study. Circle sizes are proportional to the deviation from the mean. Colours indicate the direction of such deviations: white circles represent populations showing around-average values, orange and red circles populations showing above-average values, light blue and blue circles populations showing below-average values. In each panel, the relationship between the genetic parameter and latitude is shown in the inset. Solid line represents the effect of latitude, dashed lines are the limits of 95% confidence interval. The R^2^ value of the model is also reported.

The population that contributed the most to the total allelic diversity was ORV (*A*_T_ = 0.618) while the one that contributed the least was FAB (*A*_T_ = -0.500) (Figure 5). When within-subpopulation (*A*_S_) and among-subpopulation (*D*_A_) components of *A*_T_ were disentangled, ORV still had the largest *A*_S_ value (0.461), while PAN had the lowest value of *A*_S_ (-0.948) but the largest value of *D*_A_ (0.593) (Figure 5). As expected considering their collinearity with population-specific *F*_ST_ and *Ar*, respectively, *D*_A_ exhibited a linear negative relationship with latitude (*lat*: F-value = 90.27, *P* = 3.038×10^-9^), while *A*_S_ a quadratic dependence on latitude (*lat*: F-value = 5.68, *P* = 0.026; *lat*^2^: F-value = 8.89, *P* = 0.007), with populations from central Italy showing the highest values (Figure S4). *A*_T_ showed no significant relationship with latitude (Figure S4).

**Figure 5.**
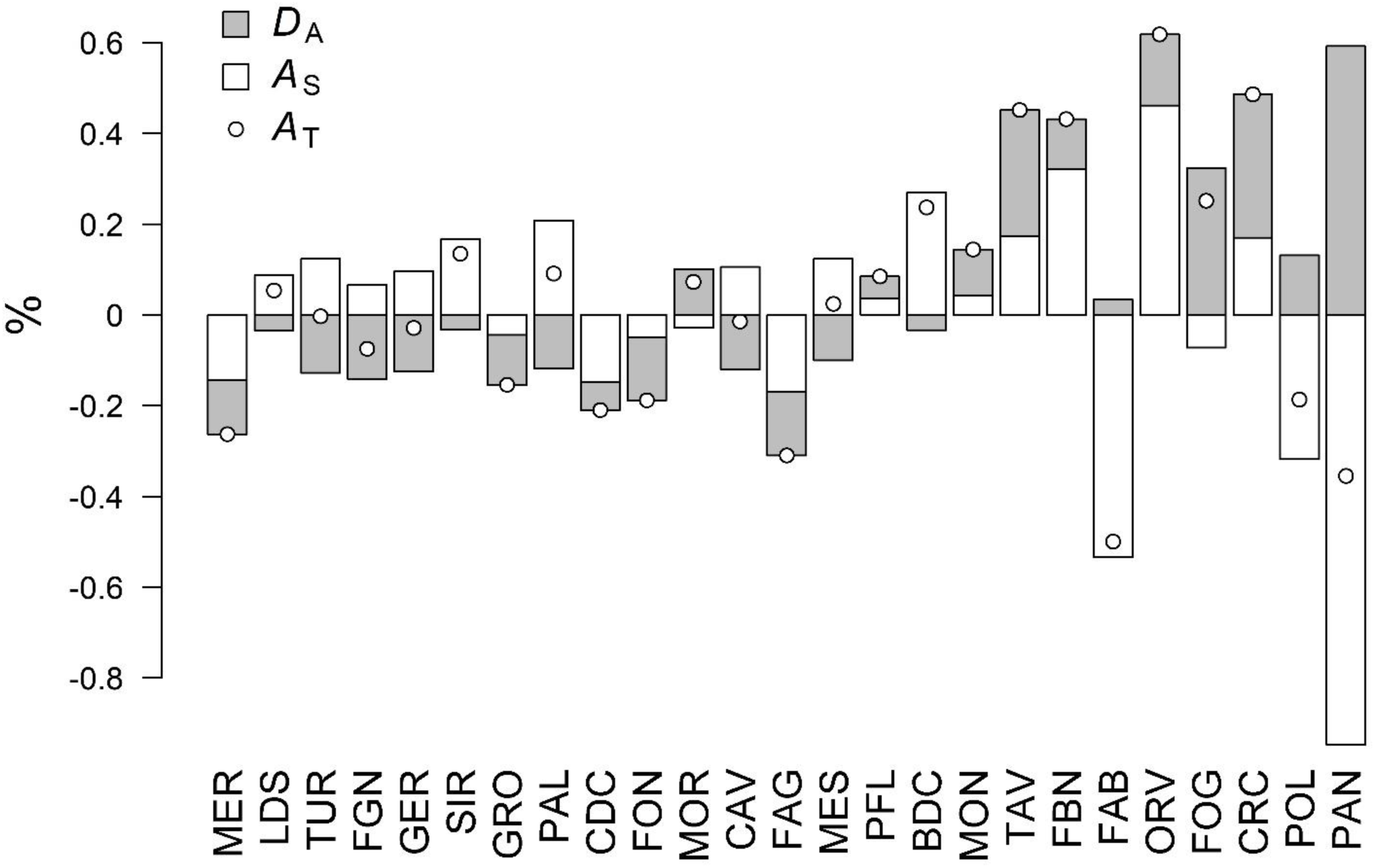
Contributions to within-(*A*_S_; white), between-(*D*_A_; grey) and total (*A*_T_; white dots) allelic diversity of *Quercus robur* populations.

### Demographic inferences

Estimates of effective population size (*Ne*) based on linkage disequilibrium did not overlap to infinite in ten cases out of 25 (Table 1). In these ten populations, mean *Ne* values were always <500 and, in three cases, <50.

No population exhibited a significant heterozygosity excess, while a significant deficiency in the MR value with respect to the values expected at the equilibrium was found in 17 populations out of 25 (Table S6), providing evidence for the presence of bottlenecks alongside most population demographic histories. In the ABC analysis, *Scenario 2* had the highest posterior probabilities (0.5918) and its confidence interval (0.5535-0.6301) did not overlap those of the other three scenarios (Table S7). Only one out of the 30 summary statistics calculated on the simulated data for *Scenario 2* was significantly different from the observed data (Table S8) which, together with the visual inspection by Principal Component Analysis (Figure S5), suggested a high goodness-of-fit of the observed data to the results of this scenario. Median values of the effective population sizes (N#) for *Scenario 2* were 6,280 (95% CI: 2,210-9710), 16,100 (95% CI:8,910-19,700), 2,440 (95% CI:711-6,650), 40,100 (95% CI: 1,990-96,600), and 6,340 (95% CI: 522-18,500) for NI, CI, SI, Nb and Na, respectively (Figure 2b, Table S9). Posterior parameters showed that CI had a larger effective population size compared to NI and SI, even considering the CI values. Median values of t1, t4, t5 and t6 were 111 (95% CI: 21-369), 671 (95% CI:564-2,070), 2,420 (95% CI: 637-7,320), and 5,220 (95% CI:1,380-9,670) generations ago, respectively (Figure 2b, Table S9). Assuming 30 years as generation time (Adams 2003), ‘gene flow’ from CI to NI (t1) and from SI to CI (t4) were dated to 3,330 (95% CI: 630-11,070) and 20,130 (95% CI: 5,280-62,100) years ago, respectively, while divergence times of NI and CI (t5) and of SI and Nb (t6) were scaled to 72,600 (95% CI: 19,110-219,600) and 156,600 (95% CI: 41,400-290,100) years ago, respectively.

Median values of ra and rd were 0.898 (95% CI: 0.429-0.993) and 0.150 (95% CI: 0.006-0.758), respectively (Figure 2b, Table S9). The median values of the mean mutation rate of SSR and mean SNI were 2.94×10^-4^ (95% CI: 1.27×10^-4^-6.74×10^-4^) and 1.67×10^-7^ (95% CI: 1.16×10^-8^-2.69×10^-6^), respectively (Table S9). The median value of the mean P was 0.280 (95% CI: 0.159-0.300) (Table S9).

## Discussion

*Quercus robur* is a keystone species in lowland ecosystems and it provides a wide range of ecosystem services and wood products. Nonetheless, there is limited knowledge on the genetic diversity of its rear edge populations, despite they reasonably harbour unique genetic variation as a legacy of southward retreatment during the Quaternary glaciations. Our study represents the first, comprehensive characterisation of the nuclear genetic diversity of *Q. robur* genetic resources across the Italian peninsula. We exhaustively covered the entire distribution of the species, sampling 745 trees from 25 populations from both large and extremely small remnant forests. As Fineschi et al. (2002), we observed the highest genetic diversity in central Italy and the lowest in southern Italy. Beyond that, we found that genetic distinctiveness and the contribution to genetic differentiation showed a south-to-north decrease. We observed a similar trend in the complexity of the genetic structure, with peninsular Italy being characterised by a mosaic of different gene pools, in contrast to the relative genetic homogeneity of populations located in northern Italy.

Multi-refugial areas are expected to host forest tree populations with more complex genetic structures than recolonization areas (de Lafontaine et al. 2013), and growing evidence is showing that such pattern holds true for several species in the Italian peninsula (e.g. Piotti et al. 2017; De Dato et al. 2020; Fortini et al. 2023). Although our results were limited to a single multi-refugial area, they showed interesting hints about pedunculate oak dynamics in the last 300,000 years. At the highest hierarchical level, both Bayesian clustering and multivariate analysis revealed the existence of two main gene pools among Italian populations: a first one, at the very rear edge of the species distribution, consisting of only one population (Bosco Pantano, PAN, along the Ionian coast), and a second one made up of all remaining populations that clearly diverged from PAN likely before the Last Glacial Maximum (LGM), even when considering the 95% credible intervals. Even though we need to be aware of the several sources of uncertainty in ABC inferences (Tsuda et al. 2015), this result points towards PAN representing the last remnant of larger southern populations that likely survived (at least) the last two glacial periods in southern Italy, isolated from the other detected gene pools. In fact, since its divergence the southern gene pool had apparently no longer received any gene flow from the north while asymmetrically acting as a source of northward historical gene flow, more intensively at the end of the LGM. Palynological and genetic evidence support the existence of a glacial refugium for pedunculate oak in the surroundings of Laghi di Monticchio, 200 km north of PAN (Watts 1996), or further south (Brewer et al. 2002; Fineschi et al. 2002; Petit et al. 2002a; Magri et al. 2015), an area for which, unfortunately, palynological records are scarce (see, for instance, Figure 1 in Magri et al. 2015). This prolonged history of isolation made PAN the most genetically distinct population in Italy, with the highest contribution to genetic differentiation and the second largest private allelic richness value. Nowadays, PAN is extremely small (i.e. 64 remaining trees, all surveyed and genotyped) and isolated, mostly surrounded by cultivated fields and coastal pine forests (Travascia et al. 2022, Pericolo et al. 2023). The population, which has undergone a strong reduction in its census size over the last century due to massive human overexploitation, showed reduced genetic variation, with an average ∼25% decrease in allelic richness with respect to the other studied populations. PAN also showed a significantly lower ratio between the number of alleles and the range of allele size than expected at equilibrium, which is usually interpreted as a sign of a severe and prolonged bottleneck (Williamson-Natesan 2005), likely eroding the otherwise high genetic diversity expected in a refugial area (Petit et al. 2003). Its F value, an analogue to *F*_ST_ between the gene pools retrieved by STRUCTURE and the assumed common ancestral population, was 0.220 for *K* = 2, which is an independent proof that the population experienced the effects of intense genetic drift (Falush et al. 2003), as expected after a severe demographic reduction. Nevertheless, no inbreeding was observed and the effective population size, albeit low, matched the census size of the population (*Ne* = 92, CI_95%_ = 67-135), probably indicating that the local gene pool was well represented in the reproductive dynamics leading to the inspected adult generation (Sandurska et al. 2019) or that the major size reduction was relatively recent, as suggested by anecdotal historical records (e.g. Douglas 1907). In this regard, it is important to stress that the genetic effects of habitat fragmentation are often overlooked in forest trees as studies have mostly focused on pre-fragmentation cohorts (Lowe et al. 2015). A strong reduction in census size due to extensive cutting in the PAN population happened in the ‘50s (Travascia et al. 2022) and, therefore, deviations in parameters more sensible to very recent demographic dynamics could only be detected in the regeneration established in the last decades, which is virtually absent in PAN.

At the next strongest levels of structuring, the gene pool comprising all remaining *Q. robur* populations was further subdivided. At *K* = 3 and *K* = 6, one and four differentiated gene pools emerged in central Italy, respectively, while northern Italian populations kept belonging to a single, rather homogeneous gene pool. We can rule out the possibility that the populations nowadays present in central and northern Italy originated from southern Italy after the Last Glacial Maximum (LGM), which was a recolonization route previously hypothesized (Fineschi et al. 2002; Petit et al. 2002a). Moreover, central Italy was not simply a ‘transit area’ where multiple recolonization routes got in contact, but it likely acted as a mosaic of small refugia where pedunculate oak persisted during the LGM, overall preserving both high and private genetic diversity. Interestingly, most populations from central Italy have linkage disequilibrium-based *Ne* estimates <500, while the global *Ne* as estimated by ABC analysis was the highest among the three gene pools. This discrepancy is compatible with a scenario where the CI gene pool originated from a mosaic of small refugia exhibiting a higher *Ne* as compared to those of nowadays, highly fragmented, declining populations. Fortini et al. (2023) reached similar conclusions for the European white oak complex in this geographic area by studying chloroplast genetic diversity in southern Europe and reviewing palaeobotanical evidence. Contrary to first syntheses (Brewer et al., 2002), more recent palaeobotanical surveys indeed reported significant percentages of deciduous *Quercus* pollen in peninsular Italy at the onset of the Holocene, with particularly high values in central Italy (Magri et al. 2015). Although a more detailed reconstruction is hampered by difficulties in distinguishing pollen grains at the species level, palynological data indicate that deciduous *Quercus* spp. had limited migrations throughout the Holocene in Italy, and major demographic events were linked to increasing or decreasing their local density (Magri et al. 2015). Our data, showing no significant clines in genetic diversity along possible recolonization routes involving specific gene pools (e.g. longitudinal clines in northern Italy), supported this hypothesis.

Northern populations showed an overall low genetic distinctiveness (average *F*_ST_ = 0.015), clustered tightly on the first three PCo axes, and showed very similar coancestry coefficients for all relevant levels of structuring in the Bayesian clustering analysis. These results were in contrast with those from a regional-scale study which reported one order of magnitude higher differentiation values (*F*_ST_ = 0.107) for *Q. robur* populations in the Po Valley based on isozyme markers (Ducci et al. 2007). However, the low levels of genetic differentiation we found in this extensive area (∼50,000 km^2^) were in line with several other studies reporting low to moderate levels of genetic differentiation at the regional scale (e.g. *G*_ST_ = 0.003 in Burger et al. 2021; *F*_ST_ = 0.032 in Kesić et al. 2021; *F*_ST_ = 0.051 in Ballian et al. 2010). *Q. robur* is indeed a wind-pollinated tree, with a high potential for long-distance dispersal and extensive gene flow (Gerber et al. 2014). Effective pollen dispersal distances up to 80 km in isolated populations are reported for this species (Buschbom et al. 2011). Altogether, available evidence about *Q. robur* gene flow capabilities and the scarce differentiation found in such a large area outlined a scenario in which *Q. robur* populations in the Po Valley constituted a single, large meta-population (*sensu* Hanski and Gilpin 1991), which has been genetically connected despite increasing fragmentation at least until the reproductive events that generated the inspected cohort. The peculiar topography of the Po Valley, with no significant geographic barriers, coupled with the current presence of typical elements of the agroforestry landscape, such as rows of *Q. robur* trees and isolated individuals, may have favoured pollen exchanges among pedunculate oak shrinking populations, leading to the maintenance of the extremely low genetic differentiation detected. Overall, our results and palaeobotanical records, showing almost complete extinction of deciduous forests during the last glacial-interglacial transition, as well as the lack of ancestral chloroplast haplotypes in northern Italy (Pini et al. 2022; Fortini et al. 2023) confirm the absence of a large refugial area in the Po plain (Petit et al., 2002a). These pieces of evidence raise the question about the post-glacial origin of these stands which currently represent by far the largest part of *Q. robur* distribution in Italy. Being highly unlikely the hypothesis of their origin from populations further south, and considering the absence of longitudinal clines in genetic diversity in this area, only enlarging the genetic characterization to populations in southern France and northern Balkans would allow to investigate a possible origin from these areas, which were classified as secondary refugia for European white oaks by Petit et al. (2002a).

The role of landscape features in shaping the genetic structure of *Q. robur* populations can be discussed also by shifting the focus on populations from central Italy, where the species occurs in much more heterogeneous topographic conditions. For instance, the distance between the two populations of ORV and FAB was ∼15 km and their pairwise genetic differentiation (*G*_ST_) was equal to 0.026. As a comparison, the easternmost and westernmost populations of the Po Valley (i.e. MER and FAG) were around 450 km apart but were much less differentiated (*G*_ST_ = 0.006). The differentiation between ORV and FAB was intercepted also by the Bayesian structuring at *K* = 6, where FAB emerged as a highly genetically peculiar population. Although the two populations were geographically close, they were separated by a small ridge that might have acted as an effective barrier to pollen and seed gene flow. As fine-scale topography may determine contrasting micro-climates across very small distance (Scherrer and Kӧrner 2011), individuals from the two populations could also experiment unsynchronised phenological cycles due to separate post-glacial dynamics, which would further hamper pollen gene flow. Despite the potential for extensive gene flow in this species, large differentiation among populations was observed at range margins (Moracho et al. 2016; Pohjanmies et al. 2016). For instance, at the southwestern margin of the species distribution, Moracho et al. (2016) studied ten *Q. robur* stands located across a 20-km long valley, identifying four distinct gene pools, and finding that most pollination events occurred within each stand. The Authors suggested that the combination of rugged topography, dense canopy and the humid microclimate of riparian forests could have limited long-distance pollen dispersal.

Although we found little genetic differentiation, especially among the northern populations, and a complete absence of inbreeding (see Table 1 and Figure S6), concerns about the genetic status of pedunculate oak populations in Italy still arise from their high to extreme degree of fragmentation. A genetic parameter which is potentially intercepting early signals of genetic decline is the effective population size (*Ne*). As it quantifies the rate at which a population loses genetic variation, *Ne* is commonly used in conservation genetics to evaluate the state and viability of populations. As a rule of thumb, a *Ne* > 50 would be beneficial for avoiding the harmful effects of inbreeding depression, while a *Ne* > 500 would be necessary for maintaining sufficient adaptive potential for coping with changing environmental conditions. (Franklin 1980). We found that ten populations had a *Ne* <500 (three of them <50). These populations represented the 25%, 62.5% and 100% of the northern, central and southern Italian populations, respectively. Low *Ne* estimates for *Q. robur* populations were often reported (Dering and Chybicki 2012; Vranckx et al. 2014; Sandurska et al. 2019). For instance, four Belgian *Q. robur* stands exhibited *Ne* estimates ranging from 23 to 58 which were associated, as in our case, to negligible inbreeding (Vranckx et al. 2014). However, these Authors found a negative relationship between *Ne* and the level of correlated paternity, suggesting that inbreeding might increase in the future because of more frequent mating among relatives. Although it is intensely debated whether *Ne* estimates could be reliable for forest tree species, particularly for large, interconnected populations (Fady and Bozzano 2021; Hoban et al. 2021; Santos-del-Blanco et al. 2022), we may consider our results on small and isolated populations, especially from central and southern Italy, as an early warning of a negative genetic trajectory that may worsen in future generations. Monitoring natural regeneration together with adult cohorts is crucial to investigate how neutral and adaptive genetic variation is transmitted through generations (Lowe et al. 2015; Vajana et al. 2023), and the low to moderate *Ne* values of our populations might predict a significant genetic depauperation in future generations.

### Conservation implications and outlook

Although post-LGM recolonization and northward expansion of tree species from Italian refugia have been widely discussed in the last two decades, this is the first study to evaluate the genetic structure and demographic history of *Quercus robur*, one of the key elements of European floodplain forests, in the Italian peninsula based on a robust sampling of extant populations. Our results showed that at least three *Q. robur* refugia contributed to the genetic layout of extant populations after the last glacial period, but refugial areas were probably even more numerous in central Italy. The complexity of the resulting genetic structure can help to identify priorities for the conservation of *Q. robur* forest genetic resources in Italy. For example, at the national scale, the identification of reserves and seed stands for in situ and ex situ conservation programs is generally not based on any knowledge on the genetic layout of forest trees’ populations. For *Q. robur*, there are almost 100 registered seed stands in Italy (MASAF 2023), the vast majority located in northern Italy (Figure 6). This means that the distinctive genetic diversity present in peninsular Italy is currently not sampled in national initiatives of seed collection despite, for instance, counting genotypes potentially pre-adapted to changing climatic conditions at northern latitudes. At the European scale, the situation improves with six Genetic Conservation Units (GCUs) of the pan-European network fostered by EUFORGEN (Koskela et al. 2013) distributed throughout the Italian range of the species (Figure 6). Three of them (corresponding to MON, CRC, and PAN) were established throughout the projects leading to the collection of the dataset presented here. However, given the mosaic of gene pools unveiled in central Italy, our results can help refine the choice of additional conservation units, possibly based on the results of algorithms of spatial conservation prioritization (Vajana et al. 2024), or ‘backup’ GCUs to further secure the genetic diversity present in each environmental zone, as recommended by EUFORGEN guidelines (de Vries et al. 2015). In addition to providing practical guidance on the conservation of genetic diversity and serving as a basis for future genetic monitoring, our results also invite for a reflection on what is the best strategy for the long-term maintenance of such small and isolated populations, that is treating them as independent conservation units or intervening to increase their genetic connectivity (e.g. through assisted migration or genetic corridors). The key to informing this choice is to integrate our results on the outcome of past demographic dynamics with an assessment of the adaptive potential and risk of non-adaptedness of *Q. robur* populations by population genomics and eco-physiological approaches in future studies.

**Figure 6.**
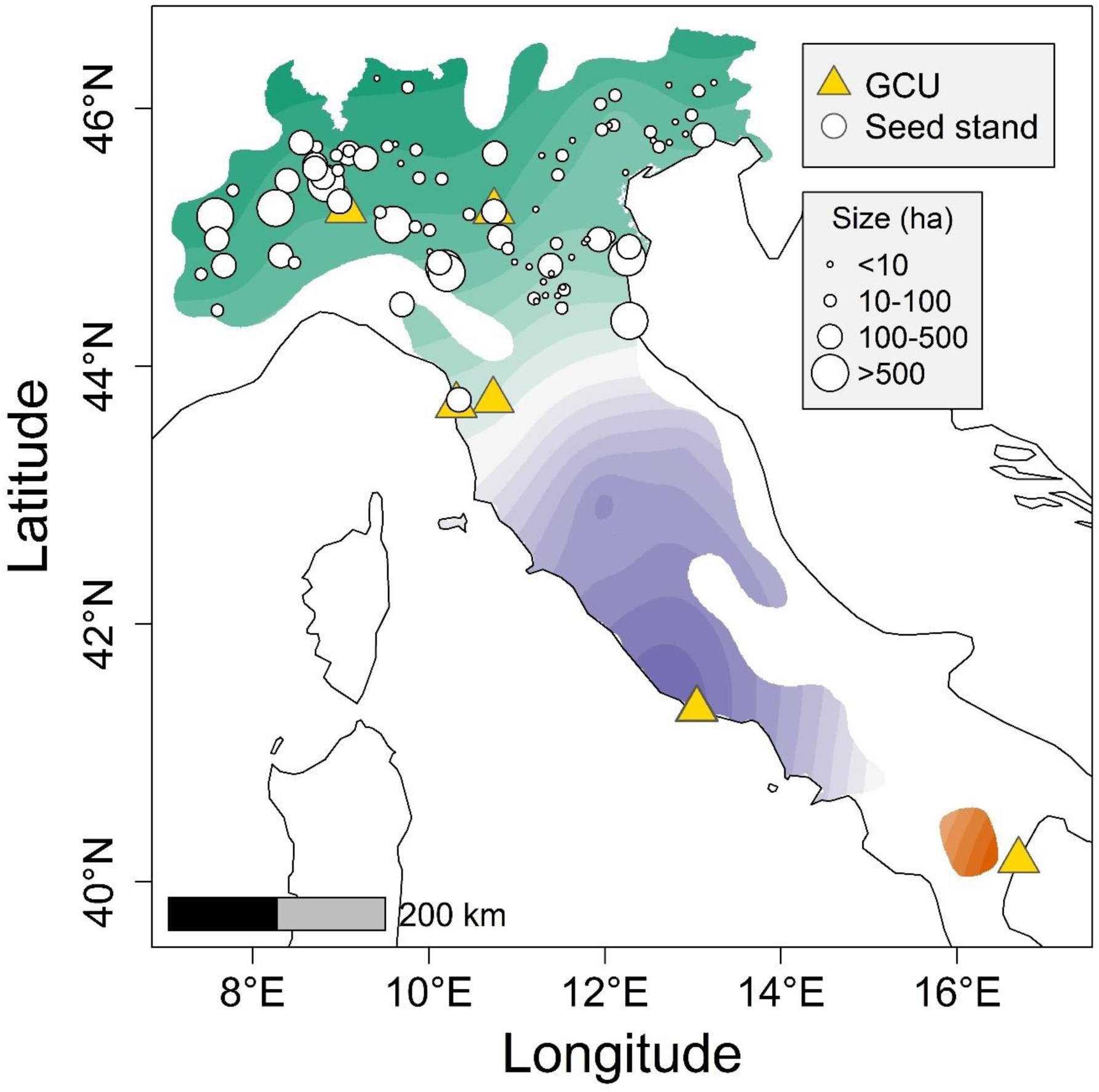
Distribution of Genetic Conservation Units (GCU) (yellow triangles) and registered seed stands (white circles) throughout the Italian peninsula. Information on the distribution of GCUs and seed stands were retrieved from EUFGIS portal (htpp://eufgis.org/) and MASAF (2023), respectively. White circles are proportional to the area of the seed stand (seed stands with an area lower than 1 ha were excluded from the representation). In the background, a spatial interpolation of admixture coefficients estimated with STRUCTURE at *K* = 3 is shown. The colour intensity indicates the probability of each pixel to belong to one of the three gene pools, based on the spatial kriging of the population q-matrix. The interpolation is limited to the distribution range of the species.

## Data Accessibility Statement

Upon acceptance of the manuscript, the dataset is privately and anonimously available for Reviewers at https://figshare.com/s/c40e3f1e9dc0d9211c0e

## APPENDIX 1

To understand which of the three populations should be set as the ancestor, we performed some pilot runs to compare three split model scenarios with northern Italy (NI), central Italy (CI) and southern Italy (SI) gene pools set as ancestral populations, respectively (Figure A). A change in population size on both branches of the tree was modelled at t1 Prior distribution of the estimated parameters are shown in Table A. The best supported scenario was Scenario A (Table B), that is the one with SI as the ancestor population, that was consequently used as ancestral population in all subsequent analyses.

**Figure A.**
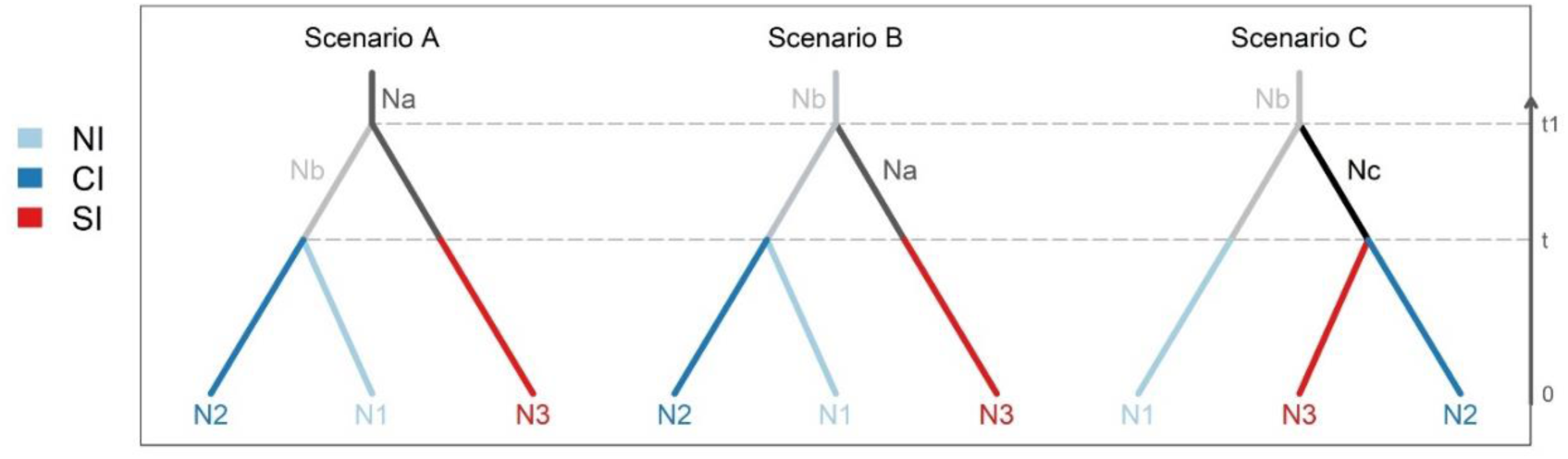
Three simple split scenarios tested to understand which populations should be set as the ancestor.

**Table A.** Prior distributions used in the DIYABC simulations.

| Parameter | Minimum | Maximum |
| --- | --- | --- |
| N1 | 10 | 10000 |
| N2 | 10 | 15000 |
| N3 | 10 | 8000 |
| Na | 10 | 20000 |
| Nb | 10 | 100000 |
| Nc | 10 | 100000 |
| $t_1$ | 10 | 10000 |
| $t$ | 10 | 1000 |
| Mean mutation rate | $1 \times 10^{-4}$ | $1 \times 10^{-3}$ |
| Individual locus mutation rate | $1 \times 10^{-5}$ | $1 \times 10^{-2}$ |
| Mean coefficient $P^{\#}$ | $1 \times 10^{-1}$ | $3 \times 10^{-1}$ |
| Individual locus coefficient $P$ | $1 \times 10^{-2}$ | $9 \times 10^{-1}$ |
| Mean SNI rate | $1 \times 10^{-8}$ | $1 \times 10^{-5}$ |
| Individual locus SNI rate | $1 \times 10^{-9}$ | $1 \times 10^{-4}$ |

**Table B.** Posterior probabilities of the three simple split scenarios tested, and their 95% confidence intervals based on the DIYABC logistic estimate.

| Scenario | Posterior probability | 95% CI (lower - upper) |
| --- | --- | --- |
| 1 | 0.9096 | 0.8925 - 0.9266 |
| 2 | 0.0599 | 0.0462 - 0.0736 |
| 3 | 0.0305 | 0.0200 - 0.0411 |

