## Supplementary Materials for "Latitudinal trends in genetic diversity and distinctiveness of *Quercus robur* rear edge forest remnants call for new conservation priorities"

**Table S1**. Geographic coordinates and sampling size of *Quercus robur* populations included in the present study.

| Population | Code | Longitude | Latitude | N sampling | N final[§] |
|---|---|---|---|---|---|
| Bosco Merlino | MER | 7.71317 | 44.78999 | 30 | 30 |
| Lame del Sesia | LDS | 8.39053 | 45.44156 | 30 | 30 |
| Turbigaccio | TUR | 8.71116 | 45.56480 | 26 | 26 |
| Fagiana | FGN | 8.82743 | 45.43451 | 30 | 30 |
| Geraci | GER | 8.98614 | 45.27786 | 30 | 30 |
| Bosco Siro Negri | SIR | 9.05683 | 45.21020 | 26[a] | 26 |
| Groane | GRO | 9.09376 | 45.66090 | 30 | 30 |
| Paladina | PAL | 9.62455 | 45.72850 | 30 | 30 |
| Capriano del Colle | CDC | 10.14840 | 45.45162 | 30[a] | 30 |
| Bosco Fontana | FON | 10.74915 | 45.19796 | 30 | 29 |
| Moriago della Battaglia | MOR | 12.13213 | 45.86176 | 30 | 30 |
| Cavalier | CAV | 12.55146 | 45.76378 | 30[a] | 30 |
| Fagagna | FAG | 13.08088 | 46.13015 | 30 | 30 |
| Bosco della Mesola | MES | 12.25539 | 44.85905 | 30 | 30 |
| Bosco della Panfilia | PFL | 11.37699 | 44.78402 | 30[a] | 30 |
| Boschi di Carrega | BDC | 10.20724 | 44.72643 | 24[a] | 24 |
| Riserva di Montefalcone | MON | 10.70773 | 43.74992 | 38 | 28 |
| Cascine di Tavola | TAV | 11.06098 | 43.84190 | 24[a] | 24 |
| Fabriano | FBN | 12.89251 | 43.35388 | 25[a] | 23 |
| Fabro | FAB | 11.98985 | 42.90642 | 24[a] | 24 |
| Orvieto | ORV | 12.09845 | 42.71113 | 24[a] | 24 |
| Foglino | FOG | 12.71482 | 41.47387 | 26[a] | 26 |
| Circeo | CRC | 13.04465 | 41.34019 | 24 | 24 |
| Pollutri | POL | 14.64420 | 42.14798 | 30 | 30 |
| Bosco Pantano | PAN | 16.68015 | 40.15543 | 64[a] | 61 |

[a] small and isolated *Q. robur* populations, with less than 500 reproductive trees

[§] final number of individuals retained for each population after excluding 'misclassified' trees based on the results of STRUCTURE

**Table S2**. Prior distributions in DIYABC simulations.

| Parameter | Minimum | Maximum |
|---|---|---|
| *Effective population size* | | |
| N1 | 10 | 10,000 |
| N2 | 10 | 20,000 |
| N3 | 10 | 8,000 |
| Na | 10 | 100,000 |
| Nb | 10 | 20,000 |
| *Time scale in generations* | | |
| t1 | 10 | 10000 |
| t2 | 10 | 10000 |
| t3 | 10 | 10000 |
| t4 | 10 | 10000 |
| t5 | 10 | 10000 |
| t6 | 10 | 10000 |
| *Admixture rates* | | |
| ra | 0.001 | 0.999 |
| rb | 0.001 | 0.999 |
| rc | 0.001 | 0.999 |
| rd | 0.001 | 0.999 |
| *Mutation model* | | |
| Mean mutation rate | $1\times10^{-4}$ | $1\times10^{-3}$ |
| Individual locus mutation rate | $1\times10^{-5}$ | $1\times10^{-2}$ |
| Mean coefficient P [#] | $1\times10^{-1}$ | $3\times10^{-1}$ |
| Individual locus coefficient P | $1\times10^{-2}$ | $9\times10^{-1}$ |
| Mean SNI rate | $1\times10^{-8}$ | $1\times10^{-5}$ |
| Individual locus SNI rate | $1\times10^{-9}$ | $1\times10^{-4}$ |

[#] parameter of the geometric distribution which is meant to generate multiple stepwise mutations

**Table S3**. Results of the ARLEQUIN analysis to detect loci under selection

| Locus | Obs. Het. BP | Obs FST | FST P-value | 1-FST quantile |
|---|---|---|---|---|
| PIE020 | 0.775185 | 0.0301432 | 0.472534 | 0.52746618 |
| PIE223 | 0.7842103 | 0.03413205 | 0.41721449 | 0.41721449 |
| PIE242 | 0.84525485 | 0.033233634 | 0.44436625 | 0.44436625 |
| PIE102 | 0.67632684 | 0.026013786 | 0.37419017 | 0.62580983 |
| PIE239 | 0.72606237 | 0.039209422 | 0.29366118 | 0.29366118 |
| PIE227 | 0.48581311 | 0.056971457 | 0.072122703 | 0.072122703 |
| PIE271 | 0.79422076 | 0.033521071 | 0.43393663 | 0.43393663 |
| PIE267 | 0.76767195 | 0.035786965 | 0.37240285 | 0.37240285 |
| PIE215 | 0.66066163 | 0.027882392 | 0.41964998 | 0.58035002 |
| QrZAG7 | 0.89793866 | 0.029775594 | 0.45564809 | 0.54435191 |
| MsQ13 | 0.76653436 | 0.0226962 | 0.27957607 | 0.72042393 |
| QrZAG112 | 0.8361915 | 0.012610696 | 0.087022963 | 0.91297704 |
| QrZAG20 | 0.84301515 | 0.036170905 | 0.36208239 | 0.36208239 |
| QpZAG15 | 0.74707049 | 0.036907397 | 0.34444895 | 0.34444895 |
| QpZAG110 | 0.57075547 | 0.033421744 | 0.45270282 | 0.45270282 |
| QrZAG96 | 0.34385221 | 0.014662097 | 0.15788062 | 0.84211938 |

**Table S4**. Results of the BayeScan analysis to detect loci under selection

| **Locus** | **Prob** | **log10(PO)** | **qVal** | **Alpha** | **Fst** |
|---|---|---|---|---|---|
| PIE020 | 0.0033 | -2.48001 | 0.887089 | -0.00082 | 0.02442 |
| PIE223 | 0.007301 | -2.1335 | 0.80298 | -0.00432 | 0.024361 |
| PIE242 | 0.004601 | -2.3352 | 0.86188 | 0.000218 | 0.024442 |
| PIE102 | 0.005801 | -2.234 | 0.83768 | -0.00086 | 0.024423 |
| PIE239 | 0.005501 | -2.2572 | 0.85075 | -0.00178 | 0.024407 |
| PIE227 | 0.0037 | -2.4301 | 0.87978 | 0.000136 | 0.024444 |
| PIE271 | 0.037004 | -1.4154 | 0.65935 | -0.03178 | 0.023905 |
| PIE267 | 0.004 | -2.3962 | 0.87146 | -0.00044 | 0.024429 |
| PIE215 | 0.006501 | -2.1842 | 0.82203 | -0.00301 | 0.02438 |
| QrZAG7 | 0.21312 | -0.56728 | 0.45845 | -0.14371 | 0.021847 |
| MsQ13 | 0.41154 | -0.1553 | 0.29423 | -0.45551 | 0.017873 |
| **QrZAG112** | **1** | **1000** | **0** | **-1.6042** | **0.005638** |
| QrZAG20 | 0.022702 | -1.634 | 0.74943 | -0.01606 | 0.024149 |
| QpZAG15 | 0.041604 | -1.3624 | 0.58343 | 0.024412 | 0.025212 |
| QpZAG110 | 0.028003 | -1.5405 | 0.71145 | -0.02095 | 0.02408 |
| QrZAG96 | 0.011901 | -1.9192 | 0.77927 | -0.00883 | 0.024293 |

**Table S5**. Matrix of pairwise $G_{ST}$. $G_{ST}$ values are below the diagonal, while their levels of significance, based on 999 permutations, are shown above the diagonal.

| | MER | LDS | TUR | FGN | GER | SIR | GRO | PAL | CDC | FON | MOR | CAV | FAG | MES | PFL | BDC | MON | TAV | FBN | FAB | ORV | FOG | CRC | POL | PAN |
|---|---|---|---|---|---|---|---|---|---|---|---|---|---|---|---|---|---|---|---|---|---|---|---|---|---|
| **MER** | | 0.107 | 0.216 | 0.109 | 0.104 | 0.048 | 0.688 | 0.037 | 0.008 | 0.037 | 0.001 | 0.170 | 0.002 | 0.001 | 0.001 | 0.027 | 0.001 | 0.001 | 0.001 | 0.001 | 0.001 | 0.001 | 0.001 | 0.001 | 0.001 |
| **LDS** | 0.002 | | 0.180 | 0.433 | 0.021 | 0.220 | 0.779 | 0.168 | 0.073 | 0.249 | 0.001 | 0.147 | 0.001 | 0.010 | 0.170 | 0.091 | 0.001 | 0.001 | 0.001 | 0.001 | 0.001 | 0.001 | 0.001 | 0.001 | 0.001 |
| **TUR** | 0.001 | 0.002 | | 0.277 | 0.015 | 0.654 | 0.417 | 0.016 | 0.001 | 0.039 | 0.001 | 0.022 | 0.001 | 0.001 | 0.014 | 0.069 | 0.007 | 0.001 | 0.001 | 0.001 | 0.001 | 0.001 | 0.001 | 0.001 | 0.001 |
| **FGN** | 0.002 | 0.000 | 0.001 | | 0.092 | 0.169 | 0.363 | 0.026 | 0.073 | 0.081 | 0.006 | 0.139 | 0.001 | 0.015 | 0.219 | 0.348 | 0.001 | 0.001 | 0.001 | 0.001 | 0.001 | 0.001 | 0.001 | 0.001 | 0.001 |
| **GER** | 0.002 | 0.004 | 0.004 | 0.002 | | 0.013 | 0.111 | 0.001 | 0.022 | 0.044 | 0.021 | 0.134 | 0.001 | 0.001 | 0.321 | 0.100 | 0.003 | 0.006 | 0.001 | 0.001 | 0.001 | 0.001 | 0.001 | 0.001 | 0.001 |
| **SIR** | 0.003 | 0.001 | -0.001 | 0.002 | 0.005 | | 0.116 | 0.115 | 0.006 | 0.038 | 0.001 | 0.020 | 0.001 | 0.012 | 0.009 | 0.016 | 0.001 | 0.001 | 0.001 | 0.001 | 0.001 | 0.001 | 0.001 | 0.001 | 0.001 |
| **GRO** | -0.001 | -0.001 | 0.000 | 0.001 | 0.002 | 0.002 | | 0.423 | 0.084 | 0.738 | 0.002 | 0.225 | 0.001 | 0.016 | 0.318 | 0.291 | 0.001 | 0.001 | 0.001 | 0.001 | 0.001 | 0.001 | 0.001 | 0.001 | 0.001 |
| **PAL** | 0.003 | 0.002 | 0.004 | 0.003 | 0.007 | 0.002 | 0.000 | | 0.069 | 0.209 | 0.001 | 0.156 | 0.001 | 0.004 | 0.005 | 0.466 | 0.001 | 0.001 | 0.001 | 0.001 | 0.001 | 0.001 | 0.001 | 0.001 | 0.001 |
| **CDC** | 0.004 | 0.003 | 0.007 | 0.003 | 0.004 | 0.006 | 0.003 | 0.003 | | 0.012 | 0.005 | 0.019 | 0.001 | 0.001 | 0.006 | 0.024 | 0.001 | 0.001 | 0.001 | 0.001 | 0.001 | 0.001 | 0.001 | 0.001 | 0.001 |
| **FON** | 0.003 | 0.001 | 0.003 | 0.003 | 0.003 | 0.003 | -0.001 | 0.001 | 0.005 | | 0.025 | 0.250 | 0.011 | 0.056 | 0.302 | 0.864 | 0.001 | 0.001 | 0.001 | 0.001 | 0.001 | 0.001 | 0.001 | 0.001 | 0.001 |
| **MOR** | 0.008 | 0.006 | 0.008 | 0.005 | 0.004 | 0.008 | 0.006 | 0.007 | 0.006 | 0.004 | | 0.315 | 0.001 | 0.012 | 0.003 | 0.004 | 0.001 | 0.001 | 0.001 | 0.001 | 0.001 | 0.001 | 0.001 | 0.001 | 0.001 |
| **CAV** | 0.001 | 0.002 | 0.004 | 0.002 | 0.002 | 0.004 | 0.001 | 0.001 | 0.004 | 0.001 | 0.001 | | 0.002 | 0.245 | 0.395 | 0.494 | 0.007 | 0.004 | 0.001 | 0.001 | 0.001 | 0.001 | 0.001 | 0.001 | 0.001 |
| **FAG** | 0.006 | 0.012 | 0.008 | 0.009 | 0.009 | 0.013 | 0.007 | 0.012 | 0.012 | 0.005 | 0.011 | 0.006 | | 0.001 | 0.001 | 0.004 | 0.001 | 0.001 | 0.001 | 0.001 | 0.001 | 0.001 | 0.001 | 0.001 | 0.001 |
| **MES** | 0.007 | 0.004 | 0.008 | 0.004 | 0.007 | 0.005 | 0.004 | 0.005 | 0.007 | 0.003 | 0.004 | 0.001 | 0.008 | | 0.034 | 0.031 | 0.001 | 0.001 | 0.001 | 0.001 | 0.001 | 0.001 | 0.001 | 0.001 | 0.001 |
| **PFL** | 0.005 | 0.001 | 0.004 | 0.001 | 0.001 | 0.004 | 0.001 | 0.006 | 0.005 | 0.001 | 0.005 | 0.000 | 0.007 | 0.003 | | 0.777 | 0.001 | 0.008 | 0.001 | 0.001 | 0.001 | 0.001 | 0.001 | 0.001 | 0.001 |
| **BDC** | 0.004 | 0.003 | 0.003 | 0.001 | 0.003 | 0.004 | 0.001 | 0.000 | 0.004 | -0.002 | 0.006 | 0.000 | 0.005 | 0.003 | -0.001 | | 0.001 | 0.001 | 0.001 | 0.001 | 0.001 | 0.001 | 0.001 | 0.001 | 0.001 |
| **MON** | 0.011 | 0.011 | 0.006 | 0.009 | 0.006 | 0.009 | 0.011 | 0.013 | 0.012 | 0.010 | 0.007 | 0.004 | 0.015 | 0.011 | 0.008 | 0.010 | | 0.001 | 0.001 | 0.001 | 0.002 | 0.001 | 0.001 | 0.001 | 0.001 |
| **TAV** | 0.011 | 0.009 | 0.011 | 0.009 | 0.007 | 0.008 | 0.009 | 0.015 | 0.012 | 0.008 | 0.008 | 0.006 | 0.015 | 0.009 | 0.005 | 0.007 | 0.007 | | 0.001 | 0.001 | 0.001 | 0.001 | 0.001 | 0.001 | 0.001 |
| **FBN** | 0.015 | 0.016 | 0.012 | 0.013 | 0.011 | 0.017 | 0.014 | 0.018 | 0.019 | 0.013 | 0.012 | 0.010 | 0.014 | 0.013 | 0.010 | 0.009 | 0.017 | 0.017 | | 0.001 | 0.001 | 0.001 | 0.001 | 0.001 | 0.001 |
| **FAB** | 0.025 | 0.025 | 0.030 | 0.029 | 0.019 | 0.032 | 0.026 | 0.029 | 0.025 | 0.028 | 0.026 | 0.023 | 0.031 | 0.031 | 0.023 | 0.027 | 0.028 | 0.031 | 0.019 | | 0.001 | 0.001 | 0.001 | 0.001 | 0.001 |
| **ORV** | 0.017 | 0.017 | 0.012 | 0.014 | 0.011 | 0.016 | 0.016 | 0.017 | 0.017 | 0.012 | 0.011 | 0.010 | 0.018 | 0.014 | 0.008 | 0.007 | 0.009 | 0.011 | 0.009 | 0.026 | | 0.001 | 0.006 | 0.001 | 0.001 |
| **FOG** | 0.028 | 0.025 | 0.020 | 0.024 | 0.020 | 0.024 | 0.025 | 0.023 | 0.027 | 0.021 | 0.020 | 0.017 | 0.025 | 0.027 | 0.015 | 0.014 | 0.023 | 0.023 | 0.015 | 0.027 | 0.009 | | 0.001 | 0.001 | 0.001 |
| **CRC** | 0.022 | 0.020 | 0.018 | 0.017 | 0.018 | 0.023 | 0.020 | 0.022 | 0.022 | 0.020 | 0.014 | 0.015 | 0.025 | 0.018 | 0.015 | 0.015 | 0.018 | 0.020 | 0.011 | 0.026 | 0.006 | 0.010 | | 0.001 | 0.001 |
| **POL** | 0.025 | 0.020 | 0.022 | 0.021 | 0.019 | 0.023 | 0.020 | 0.023 | 0.026 | 0.017 | 0.023 | 0.016 | 0.028 | 0.018 | 0.021 | 0.017 | 0.026 | 0.025 | 0.016 | 0.034 | 0.022 | 0.024 | 0.016 | | 0.001 |
| **PAN** | 0.045 | 0.048 | 0.047 | 0.044 | 0.042 | 0.052 | 0.045 | 0.048 | 0.054 | 0.046 | 0.048 | 0.039 | 0.043 | 0.047 | 0.041 | 0.036 | 0.045 | 0.048 | 0.034 | 0.053 | 0.035 | 0.039 | 0.034 | 0.037 | |

**Table S6**. Heterozygosity excess and M-ratio deficiency tests for assessing the occurrence of past genetic bottlenecks in *Quercus robur* populations. *H* and MR are, respectively, the observed values of expected heterozygosity and the M-ratio, while Δ*H* and ΔMR are the differences between such observed values and the values expected at the equilibrium. *P* is the *P*-value of the Wilcoxon signed-rank test used to evaluate the statistical significance of such differences (° *P*<0.1, * *P*<0.05, ** *P*<0.01, *** *P*<0.001).

| **Population** | ***H*** | **Δ*H*** | ***P*** | | **MR** | **ΔMR** | ***P*** |
|---|---|---|---|---|---|---|---|
| MER | 0.6895 | 0.0517 | 0.6833 | | 0.6960 | 0.0913 | 0.0223* |
| LDS | 0.7104 | 0.0601 | 0.9529 | | 0.7133 | 0.0623 | 0.0889° |
| TUR | 0.7051 | 0.0662 | 0.9627 | | 0.6597 | 0.1102 | 0.0526° |
| FGN | 0.7016 | 0.0642 | 0.8840 | | 0.7357 | 0.0411 | 0.3920 |
| GER | 0.6866 | 0.0796 | 0.9711 | | 0.6639 | 0.1128 | 0.0031** |
| SIR | 0.7407 | 0.0268 | 0.6278 | | 0.6884 | 0.0821 | 0.0373* |
| GRO | 0.6958 | 0.0642 | 0.9416 | | 0.6679 | 0.1128 | 0.0077** |
| PAL | 0.7088 | 0.0625 | 0.9414 | | 0.7299 | 0.0449 | 0.2176 |
| CDC | 0.6957 | 0.0476 | 0.7195 | | 0.6997 | 0.0861 | 0.0370* |
| FON | 0.6854 | 0.0686 | 0.9128 | | 0.7021 | 0.0787 | 0.1494 |
| MOR | 0.6940 | 0.0681 | 0.9631 | | 0.6656 | 0.1138 | 0.0469* |
| CAV | 0.7085 | 0.0555 | 0.9280 | | 0.6930 | 0.0856 | 0.0531° |
| FAG | 0.6649 | 0.0778 | 0.9955 | | 0.6829 | 0.1045 | 0.0368* |
| MES | 0.7184 | 0.0585 | 0.9585 | | 0.6961 | 0.0769 | 0.0657° |
| PFL | 0.7059 | 0.0485 | 0.8739 | | 0.6469 | 0.1335 | 0.0091** |
| BDC | 0.7107 | 0.0668 | 0.9035 | | 0.7023 | 0.0603 | 0.1383 |
| MON | 0.6968 | 0.0748 | 0.9935 | | 0.7037 | 0.0695 | 0.0796° |
| TAV | 0.7152 | 0.0575 | 0.9631 | | 0.7061 | 0.0604 | 0.2182 |
| FBN | 0.7175 | 0.0466 | 0.9124 | | 0.7333 | 0.0310 | 0.2989 |
| FAB | 0.6369 | 0.0643 | 0.9478 | | 0.6801 | 0.1178 | 0.0531° |
| ORV | 0.7410 | 0.0534 | 0.9962 | | 0.7643 | -0.0091 | 0.4931 |
| FOG | 0.7050 | 0.0384 | 0.9283 | | 0.6830 | 0.0977 | 0.0195* |
| CRC | 0.7212 | 0.0416 | 0.9537 | | 0.7577 | 0.011 | 0.5511 |
| POL | 0.6777 | 0.0482 | 0.8391 | | 0.7023 | 0.0902 | 0.0528° |
| PAN | 0.6562 | 0.0072 | 0.5897 | | 0.6378 | 0.1913 | 0.0002*** |

**Table S7**. Posterior probabilities of the four demographic scenarios tested and their 95% confidence intervals based on DIYABC logistic estimation.

| **Scenario** | **Posterior probability** | **95% CI (lower - upper)** |
|---|---|---|
| 1 | 0.2205 | 0.1910 - 0.2500 |
| 2 | 0.5918 | 0.5535 - 0.6301 |
| 3 | 0.1582 | 0.1310 - 0.1848 |
| 4 | 0.0295 | 0.0182 - 0.0408 |

**Table S8**. Model checking for *Scenario 2*. Comparison of summary statistics for the observed data sets and simulated posterior data sets.

| **Summary statistics** | **Observed value** | ***Scenario 2* proportion (simulated<observed)** |
|---|---|---|
| Mean number of alleles NI | 10.250 | 0.5040 |
| Mean number of alleles CI | 10.500 | 0.5935 |
| Mean number of alleles SI | 6.375 | 0.5405 |
| Mean expected heterozygosity NI | 0.712 | 0.2705 |
| Mean expected heterozygosity CI | 0.719 | 0.4040 |
| Mean expected heterozygosity SI | 0.656 | 0.6180 |
| Mean allele size variance NI | 10.342 | 0.4950 |
| Mean allele size variance CI | 7.831 | 0.3690 |
| Mean allele size variance SI | 6.141 | 0.4335 |
| Mean Garza-Williamson's M NI | 0.713 | 0.1070 |
| Mean Garza-Williamson's M CI | 0.785 | 0.3515 |
| Mean Garza-Williamson's M SI | 0.597 | 0.1010 |
| Mean number of alleles NI&CI | 12.313 | 0.5865 |
| Mean number of alleles NI&SI | 11.188 | 0.5375 |
| Mean number of alleles CI&SI | 11.375 | 0.6240 |
| Mean expected heterozygosity NI&CI | 0.725 | 0.3715 |
| Mean expected heterozygosity NI&SI | 0.713 | 0.4635 |
| Mean expected heterozygosity CI&SI | 0.710 | 0.5140 |
| Mean allele size variance NI&CI | 9.182 | 0.4340 |
| Mean allele size variance NI&SI | 8.768 | 0.4720 |
| Mean allele size variance CI&SI | 7.206 | 0.3820 |
| $F_{ST}$ NI&CI | 0.026 | 0.9540* |
| $F_{ST}$ NI&SI | 0.078 | 0.5850 |
| $F_{ST}$ CI&SI | 0.063 | 0.4925 |
| Mean individual assignment likelihood NI&CI | 1.427 | 0.3625 |
| Mean individual assignment likelihood NI&SI | 2.101 | 0.4875 |
| Mean individual assignment likelihood CI&NI | 1.458 | 0.5310 |
| Mean individual assignment likelihood CI&SI | 1.955 | 0.5125 |
| Mean individual assignment likelihood SI&NI | 1.569 | 0.6685 |
| Mean individual assignment likelihood SI&CI | 1.528 | 0.6910 |

* Proportions lower than 5% or greater than 95%

**Table S9**. Parameter estimates for *Scenario 2* which was the best demographic scenario according to posterior probabilities.

| | | | | Quantiles | | | |
|---|---|---|---|---|---|---|---|
| **Parameter** | **Mean** | **Median** | **Mode** | **2.5%** | **5%** | **95%** | **97.5%** |
| N1 | 6.22e+003 | 6.28e+003 | 6.30e+003 | 2.21e+003 | 2.73e+003 | 9.46e+003 | 9.71e+003 |
| N2 | 1.56e+004 | 1.61e+004 | 1.71e+004 | 8.91e+004 | 1.02e+004 | 1.94e+004 | 1.97e+004 |
| N3 | 2.75e+003 | 2.44e+003 | 1.99e+003 | 7.11e+002 | 9.00e+002 | 5.88e+003 | 6.65e+003 |
| Na | 7.41e+003 | 6.34e+003 | 3.68e+003 | 5.22e+002 | 8.89e+002 | 1.72e+004 | 1.85e+004 |
| Nb | 4.36e+004 | 4.01e+004 | 4.09e+003 | 1.99e+003 | 3.82e+003 | 9.30e+004 | 9.66e+004 |
| t1 | 1.32e+002 | 1.11e+002 | 8.28e+001 | 2.10e+001 | 2.85e+001 | 3.07e+002 | 3.69e+002 |
| t4 | 7.81e+002 | 6.71e+002 | 5.64e+002 | 1.76e+002 | 2.32e+002 | 1.71e+003 | 2.07e+003 |
| t5 | 2.86e+003 | 2.42e+003 | 1.75e+003 | 6.37e+002 | 7.81e+002 | 6.44e+003 | 7.32e+003 |
| t6 | 5.38e+003 | 5.22e+003 | 3.99e+003 | 1.38e+003 | 1.70e+003 | 9.40e+003 | 9.67e+003 |
| ra | 8.56e-001 | 8.98e-001 | 9.44e-001 | 4.29e-003 | 5.82e-001 | 9.88e-001 | 9.93e-001 |
| rd | 2.04e-001 | 1.50e-001 | 1.00e-003 | 6.47e-003 | 1.31e-002 | 6.16e-001 | 7.58e-001 |
| Mean mutation rate | 3.22e-004 | 2.94e-004 | 2.49e-004 | 1.27e-004 | 1.42e-004 | 6.00e-004 | 6.74e-004 |
| Mean coefficient P | 2.67e-001 | 2.80e-001 | 3.00e-001 | 1.59e-001 | 1.82e-001 | 3.00e-001 | 3.00e-001 |
| Mean SNI rate | 4.75e-007 | 1.67e-007 | 1.09e-008 | 1.16e-008 | 1.35e-008 | 2.01e-006 | 2.69e-006 |

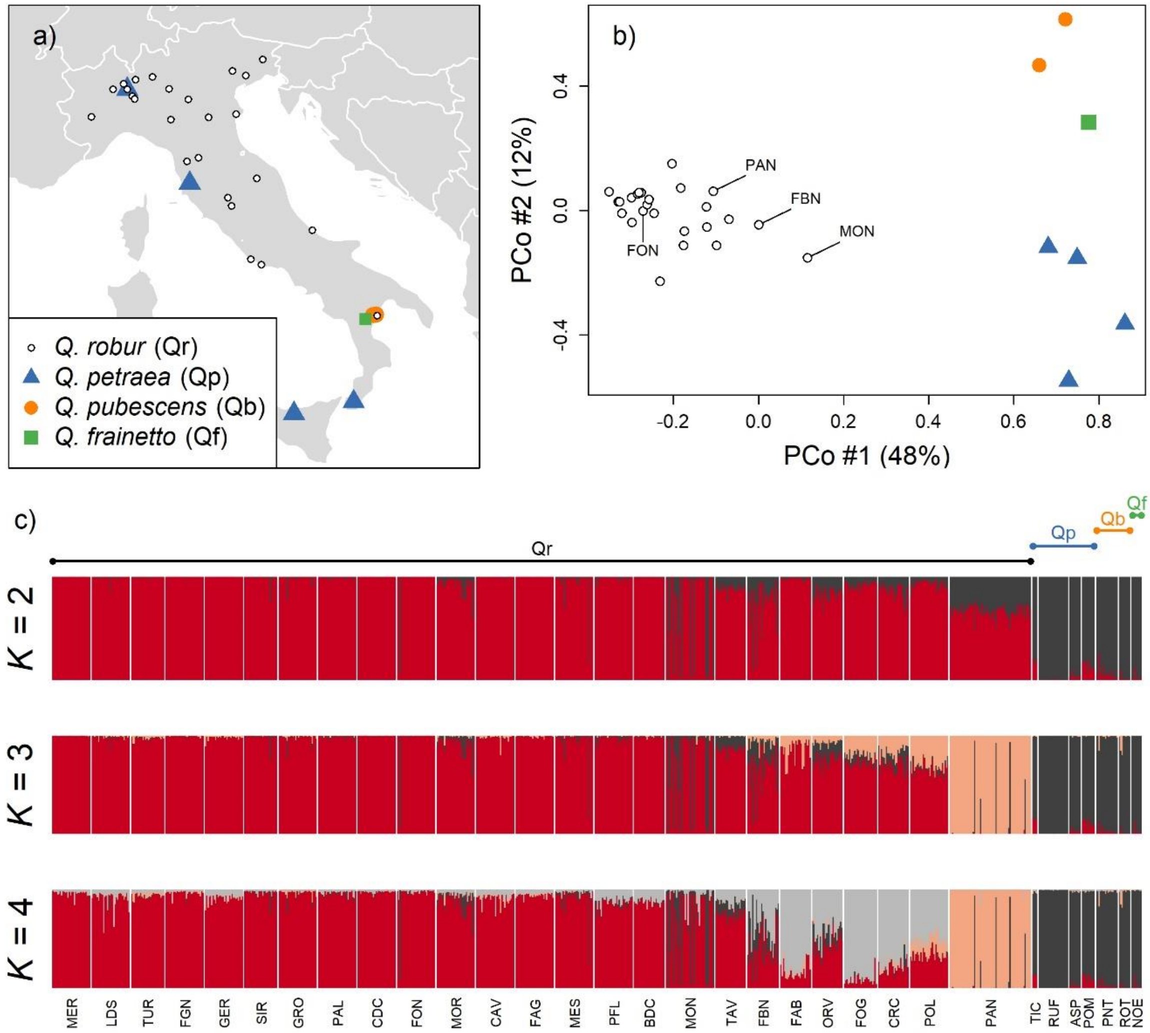

**Figure S1**. (**a**) Map of white oak populations used to identify 'misclassified' trees. (**b**) Results of a Principal Coordinate Analysis made on mean among-population genetic distances. Populations in which 'misclassified' individuals have been found are highlighted in the graph with their label. (**c**) Results of the Bayesian clustering analysis run with STRUCTURE. Results are shown for *K* = 2, *K* = 3 and *K* = 4.

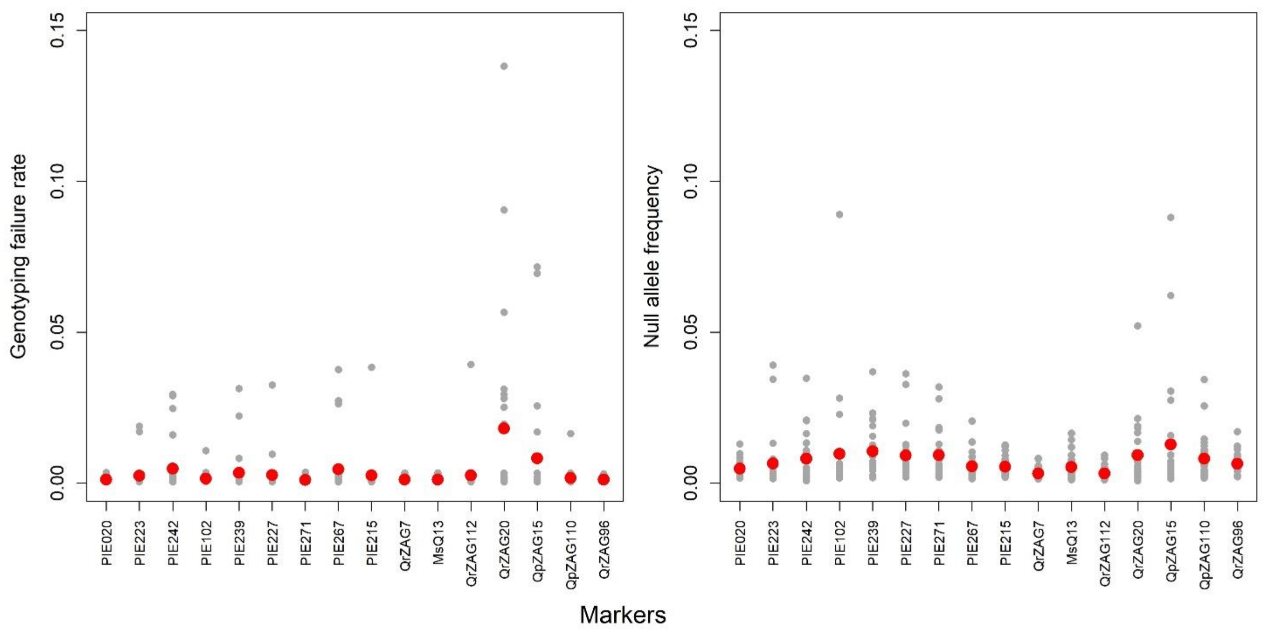

**Figure S2**. Mean values of genotyping failure rates and null allele frequencies estimated for population and microsatellite locus by using INest. Red dots represent genotyping failure rates and null allele frequencies averaged over populations.

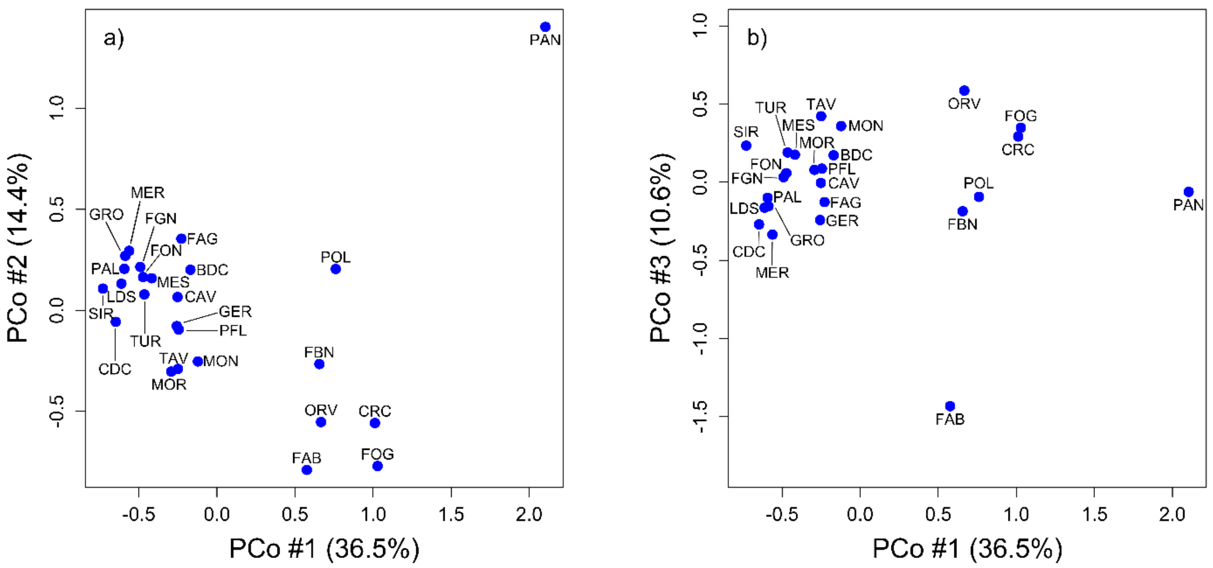

**Figure S3**. Principal Coordinate Analysis (PCoA) on the matrix of pairwise genetic distances among *Quercus robur* populations. First *vs*. second Principal Coordinate (PCo) scores are shown on the left-side panel, while first *vs*. third PCo scores are shown on the right-side panel.

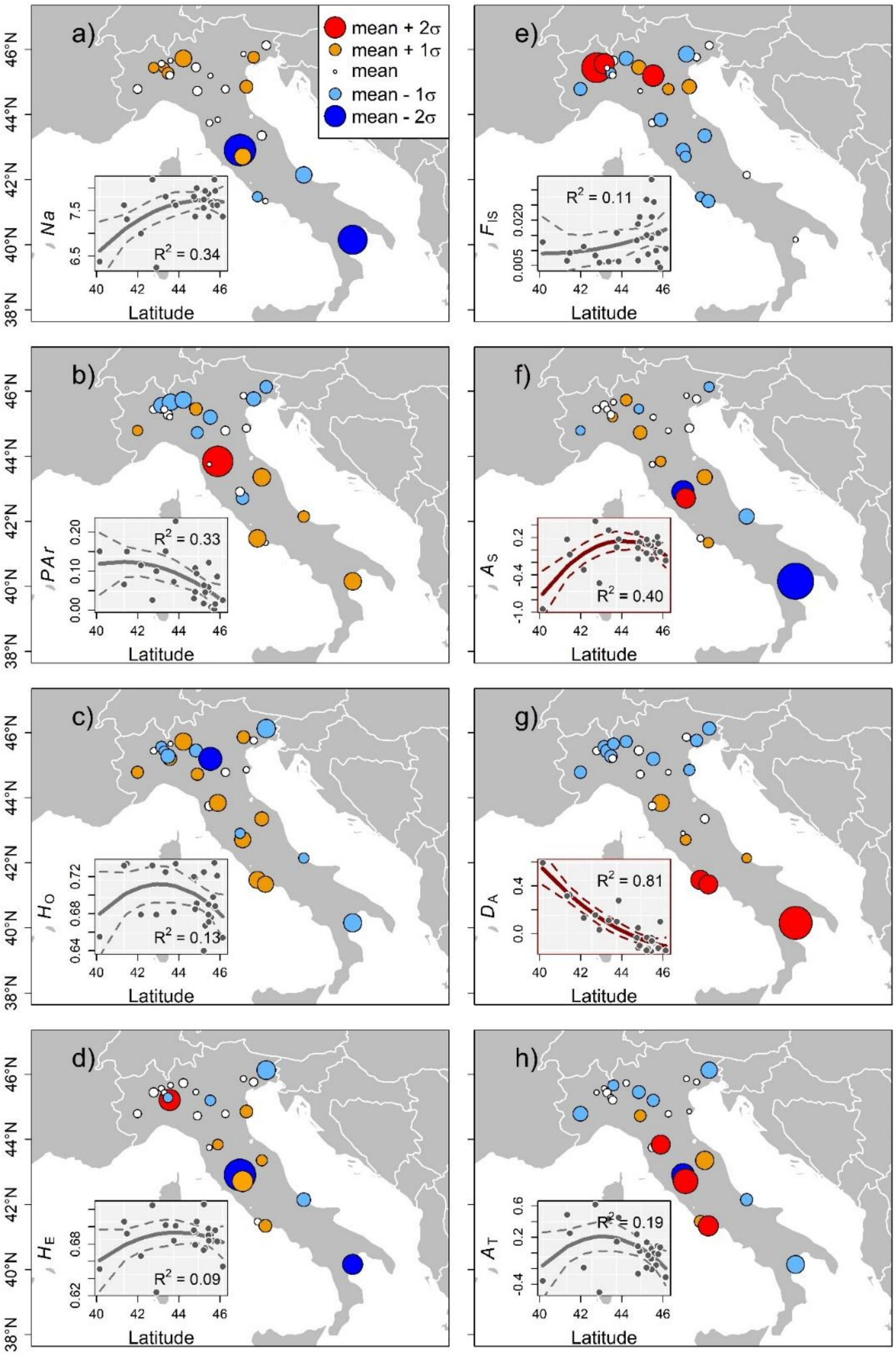

**Figure S4**. Estimates of (a) number of alleles (*Ar*), (b) private allelic richness (*PAr*), (c) observed heterozygosity ($H_O$), (d) expected heterozygosity ($H_E$), (e) inbreeding coefficient ($F_{IS}$), and (f) contribution to within-population ($A_S$), (g) between-population ($D_A$) and (i) total allelic diversity ($A_T$) in the 25 *Quercus robur* populations of the present study. Circle sizes are proportional to the deviation from the mean. Colours indicate the direction of such deviations: white circles represent populations showing around-average values, orange and red circles populations showing above-average values, light blue and blue circles populations showing below-average values. In each panel, the relationship between genetic parameter and latitude is shown in the inset. When latitude has a statistically significant effect on the genetic parameter, curves are coloured in darkred, otherwise in grey. The $R^2$ value of the model is also reported.

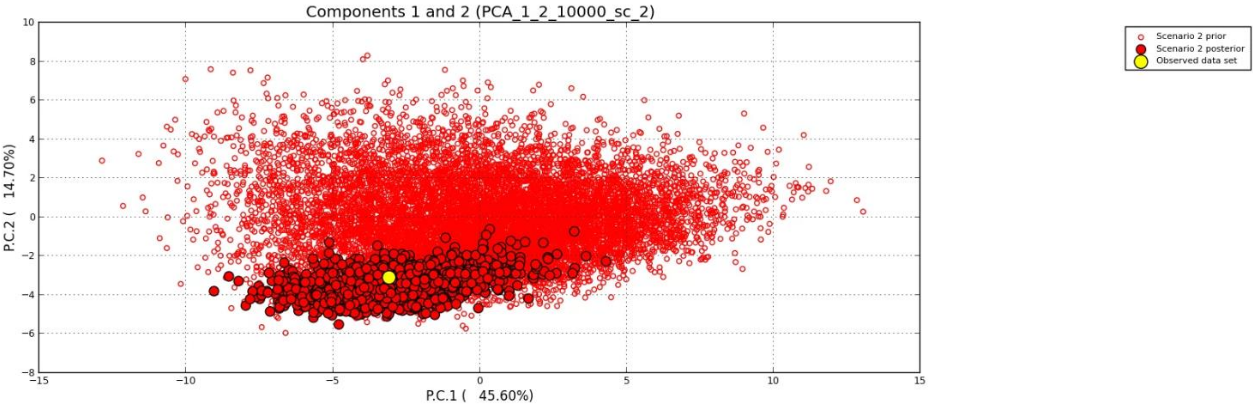

**Figure S5**. Principal Component Analysis (PCA) obtained by DIYABC for *Scenario 2.*

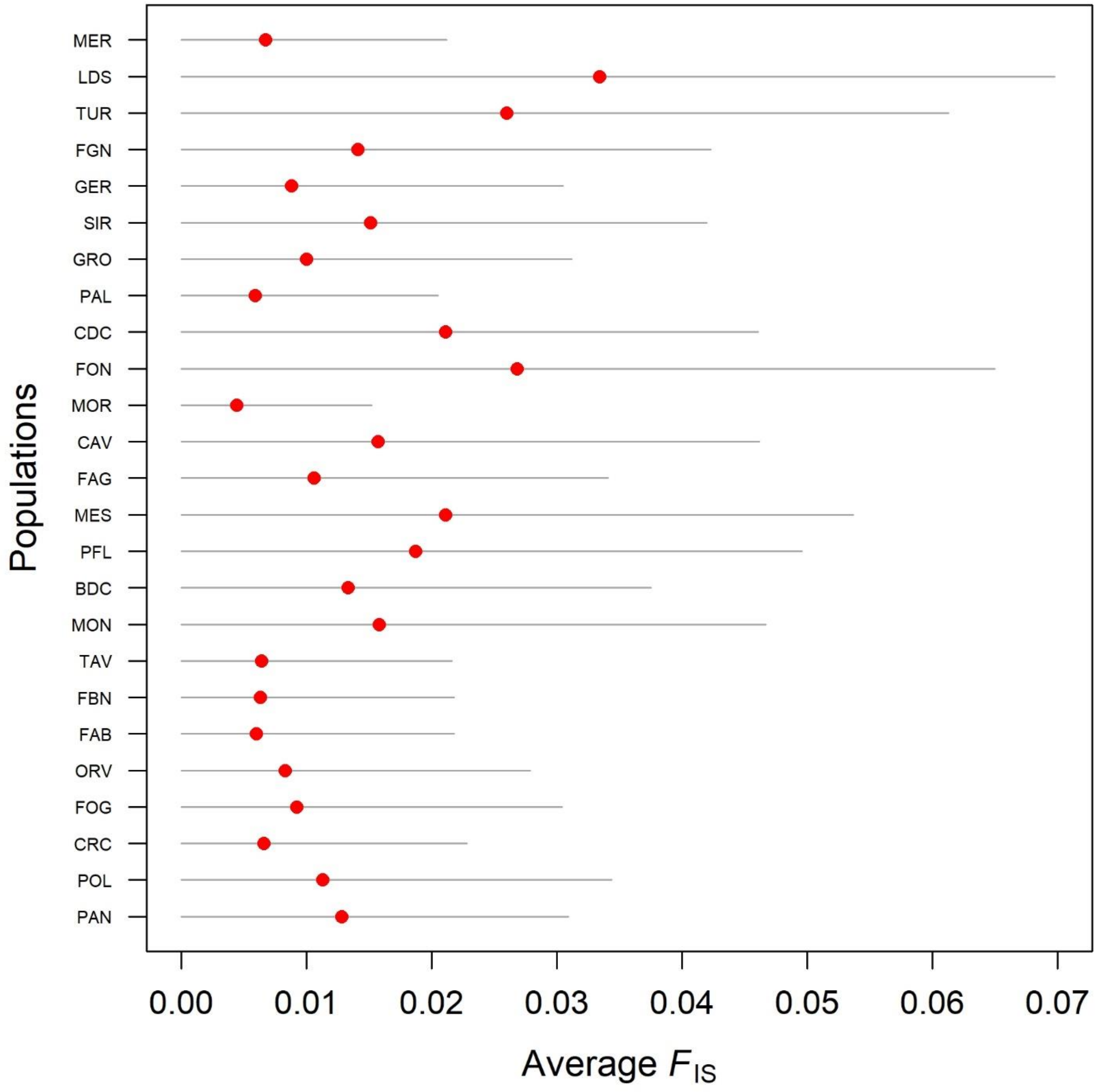

**Figure S6**. Average inbreeding coefficient ($F_{IS}$) estimated for each population by using INest. Grey lines represent the 95% highest density intervals of $F_{IS}$ posterior distributions.
